# Rapid heuristic inference of antibiotic resistance and susceptibility by genomic neighbor typing

**DOI:** 10.1101/403204

**Authors:** Karel Břinda, Alanna Callendrello, Kevin C. Ma, Derek R MacFadden, Themoula Charalampous, Robyn S Lee, Lauren Cowley, Crista B Wadsworth, Yonatan H Grad, Gregory Kucherov, Justin O’Grady, Michael Baym, William P Hanage

## Abstract

Surveillance of drug-resistant bacteria is essential for healthcare providers to deliver effective empiric antibiotic therapy. However, traditional molecular epidemiology does not typically occur on a timescale that could impact patient treatment and outcomes. Here we present a method called ‘genomic neighbor typing’ for inferring the phenotype of a bacterial sample by identifying its closest relatives in a database of genomes with metadata. We show that this technique can infer antibiotic susceptibility and resistance for both *S. pneumoniae* and *N. gonorrhoeae*. We implemented this with rapid *k*-mer matching, which, when used on Oxford Nanopore MinION data, can run in real time. This resulted in determination of resistance within ten minutes (sens/spec 91%/100% for *S. pneumoniae* and 81%/100% *N. gonorrhoeae* from isolates with a representative database) of sequencing starting, and for clinical metagenomic sputum samples (75%/100% for *S. pneumoniae*), within four hours of sample collection. This flexible approach has wide application to pathogen surveillance and may be used to greatly accelerate appropriate empirical antibiotic treatment.

## Introduction

Infections pose multiple challenges to healthcare systems, contributing to higher mortality, morbidity, and escalating cost. Clinicians must regularly make rapid decisions on empiric antibiotic treatment of infectious syndromes without knowing the causative pathogen(s) or whether they are drug-susceptible or drug-resistant. In some cases, this is directly linked to poor outcomes; in the case of septic shock, the risk of death increases by an estimated 10% with every 60 minutes delay in initiating effective treatment^1^.

The molecular epidemiology of infectious disease allows us to identify high-risk pathogens and determine their patterns of spread, on the basis of their genetics or (increasingly) genomics. Conventionally such studies, including outbreak investigations and characterization of novel resistant strains, have been conducted in retrospect, but this has been changing with the availability of new and increasingly inexpensive sequencing technologies^2, 3^. The wealth of data generated by genomics is promising but introduces a new challenge: while many features of a sequence are correlated with the phenotype of interest, few are causative.

Prescription, however, has long been informed by correlative features when causative ones are difficult to measure, for example whether the same syndrome or pathogen occurring in other patients from the same clinical environment have responded to a particular antibiotic. This has also been observed at the genetic level as well, as a result of genetic linkage between resistance elements and the rest of the genome. An example is given by the pneumococcus (*Streptococcus pneumoniae*). The Centers for Disease Control have rated the threat level of drug-resistant pneumococcus as ‘serious’ ^4^. While resistance arises in pneumococci through a variety of mechanisms, approximately 90% of the variance in the minimal inhibitory concentration (MIC) for antibiotics of different classes can be explained by the loci determining the strain type^5^, even though none of these loci themselves causes resistance. Thus, in the overwhelming majority of cases, resistance and susceptibility can be inferred from coarse strain typing based on population structure. This population structure could be leveraged to offer an alternative approach to detecting resistance in which rather than detecting high-risk genes, we identify high-risk strains. While many approaches have been developed to identify whether a pathogen carries mutations or genes known to confer resistance^6–21^ (see ref^22^ for a comprehensive review), this is not equivalent to the clinical question of whether the pathogen is susceptible.

We present a method called ‘genomic neighbor typing’ which can bring molecular epidemiology closer to the bedside and provide information relevant to treatment at a much earlier stage. Our method takes sequences generated from a sample in ‘real time’ and matches them to a database of genomes to identify the closest relatives. Because closely related isolates usually have similar properties, this yields an informed heuristic as to the pathogen’s phenotype. We demonstrate this by identifying drug-resistant and drug-susceptible clones for both *Streptococcus pneumoniae* (the pneumococcus) and *Neisseria gonorrhoeae* (the gonococcus), within minutes after the start of sequencing using Oxford Nanopore Technology. The method has many potential applications, depending on the specific pathogen and quality of the databases available for matching, which we discuss together with its limitations.

## Results

### Resistance is associated with clones in S. pneumoniae and N. gonorrhoeae

To quantify the association of clones with antibiotic resistance of the pathogens *S. pneumoniae* and *N. gonorrhoeae*, we constructed optimal predictors of resistance from bacterial lineages and measured the associated Area under the Receiver Operation Characteristic Curve (AUC) (**Supplementary Document 1**). First, we applied the method to 616 pneumococcal genomes from a carriage study in Massachusetts children^23, 24^. Second, we used 1102 clinical gonococcal isolates collected from 2000 to 2013 by the Centers for Disease Control and Prevention’s Gonococcal Isolate Surveillance Project^25^. In both cases, the datasets comprised draft genome assemblies from Illumina HiSeq reads, resistance data, and lineages inferred from sequence cluster computed using Bayesian Analysis of Population Structure (BAPS)^26^. Lineages of *S. pneumoniae* are predictive for benzylpenicillin, ceftriaxone, trimethoprim-sulfamethoxazole, erythromycin, and tetracycline resistance with AUC ranging from 0.90 to 0.97 (**Supplementary Document 1**), consistent with previous works^5^. In *N. gonorrhoeae*, ciprofloxacin, ceftriaxone, and cefixime attained comparably large AUCs (from 0.93 to 0.98) whereas azithromycin demonstrated lower association (AUC 0.80), as observed previously^25^.

### Rapid identification of nearest known relative from sequencing reads

Based on the observed associations we developed an approach that we term ‘genomic neighbor typing’ to predict phenotype from sequencing data. Genomic neighbor typing is a two-step algorithm, which first compares a provided sample to a database of reference genomes with a known phylogeny and phenotype, and then predicts the likely phenotype of the sample based on the best hits (nearest neighbors) and their matching quality. We apply this here to the detection of drug resistance.

To implement genomic neighbor typing we developed software called RASE (Resistance-Associated Sequence Elements) (**Figure 1**). RASE takes a stream of nanopore reads and compares their *k*-mer content to references using a modified version of ProPhyle^27, 28^, a metagenomic classifier implementing a fast and memory-efficient exact colored de Bruijn graph data structure^29^ using a BWT index^30^ (Methods). Using ProPhyle RASE identifies which references are the most similar to the read and increases their similarity weights (this approach was inspired by but differs from other similar approaches such as Kraken^31^ and Kallisto^32^). These weights are cumulative scores capturing sample-to-reference similarity; they are set to zero at the beginning and are increased on-the-fly as sequencing proceeds according to each read’s ‘information content’ (Methods). Generally speaking, longer reads, such as those covering multiple accessory genes, tend to be specific and have high scores, whereas short reads or reads from the core genome are found in many lineages, tend to be non-specific and have low scores. Weights serve as a proxy to inverted genetic distance between the sample and the references.

**Figure 1:**
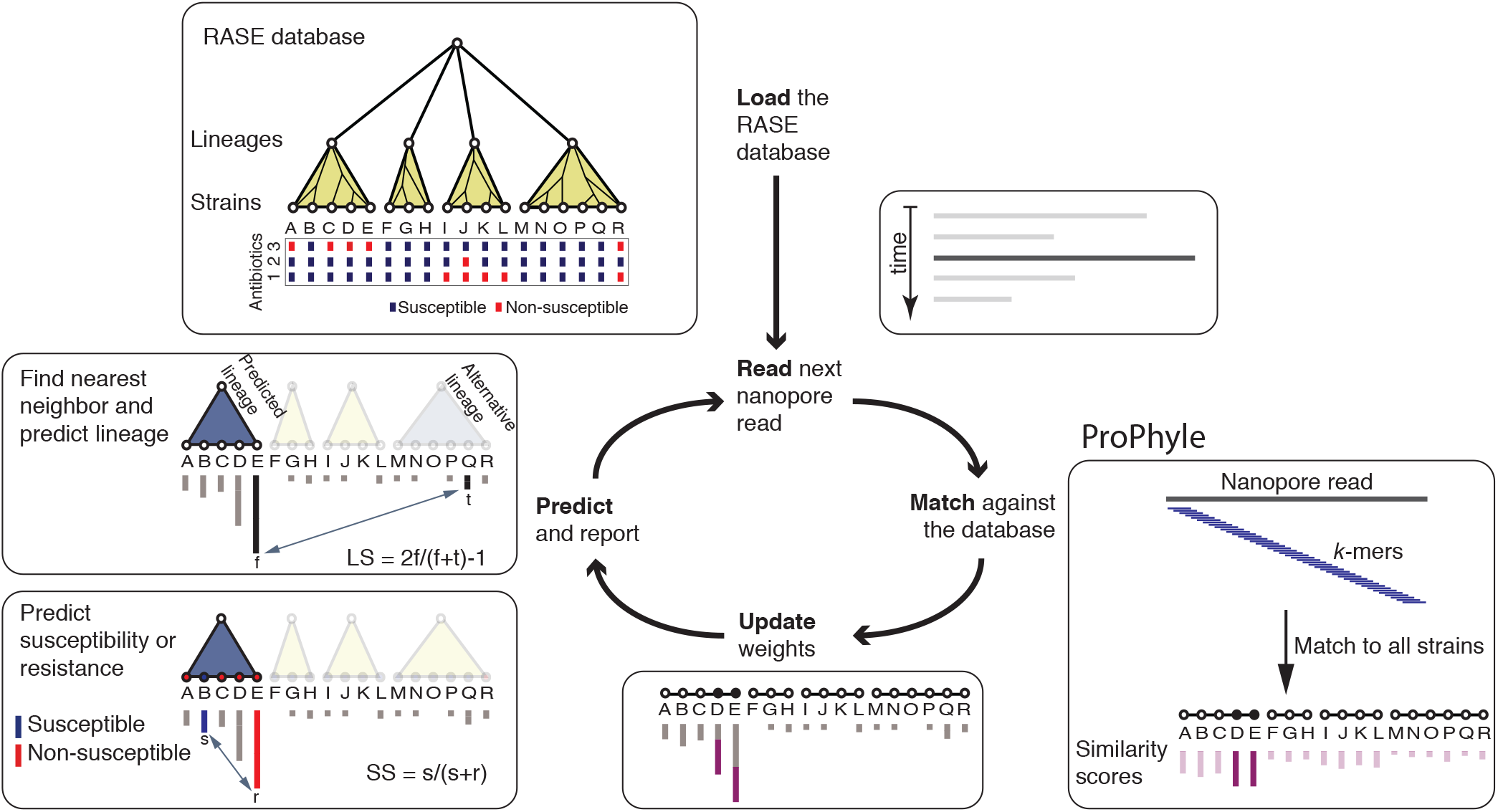
Overview of the RASE approach. In the first, loading step, the precomputed RASE database is loaded into memory. As reads are generated, they are matched against the database using ProPhyle to calculate similarity to individual strains. The weights for the most similar strains (D and E in the figure) are increased proportionate to the number of matching *k*-mers. Finally, resistance is predicted from the obtained weights and the resistance profiles of the database strains as follows: First, the best lineage is identified as the lineage of the best match (having the highest weight, E in the figure) and its score is calculated (lineage score, LS). Second, for every antibiotic, a score quantifying the chance of susceptibility (susceptibility score, SS) is calculated, based on the most similar susceptible and resistant strains inside the identified lineage (B and E in the figure, respectively). The susceptibility or resistance to each of the antibiotics is predicted from their susceptibility scores by a comparison with a threshold (0.5 in the default setting), and reported together with the lineage, the best matching strain and that strain’s known properties (e.g., the original antibiograms, MLST sequence type, or serotype).

Resistance or susceptibility is predicted in two steps based on the computed weights, the population structure, and the reference phenotypes. First, RASE identifies the lineage of the best matching reference genome and estimates the confidence of lineage assignment by comparing the two best matching lineages to compute a ‘lineage score’ (Methods). Subsequently, RASE identifies the best match within that lineage and predicts resistance from the nearest resistant and susceptible neighbors. Comparison of their weights provides a ‘susceptibility score’, which quantifies the risk of resistance (Methods). When the weights are too similar, the call’s confidence is considered low; this happens when resistant and susceptible strains are insufficiently genetically distinct, which is often the case for resistance emerging recently in evolutionary history (Methods). The ability to pinpoint the closest relatives in the database offers further resolution, even in the case where the resistance phenotype varies within a lineage.

Results of RASE are reported in real time as the best match in the database, together with susceptibility scores to the antibiotics being tested and a proportion of matching *k*-mers for quality control. As the run progresses, the scores fluctuate and eventually stabilize (an example shown in **Figure 2**).

**Figure 2:**
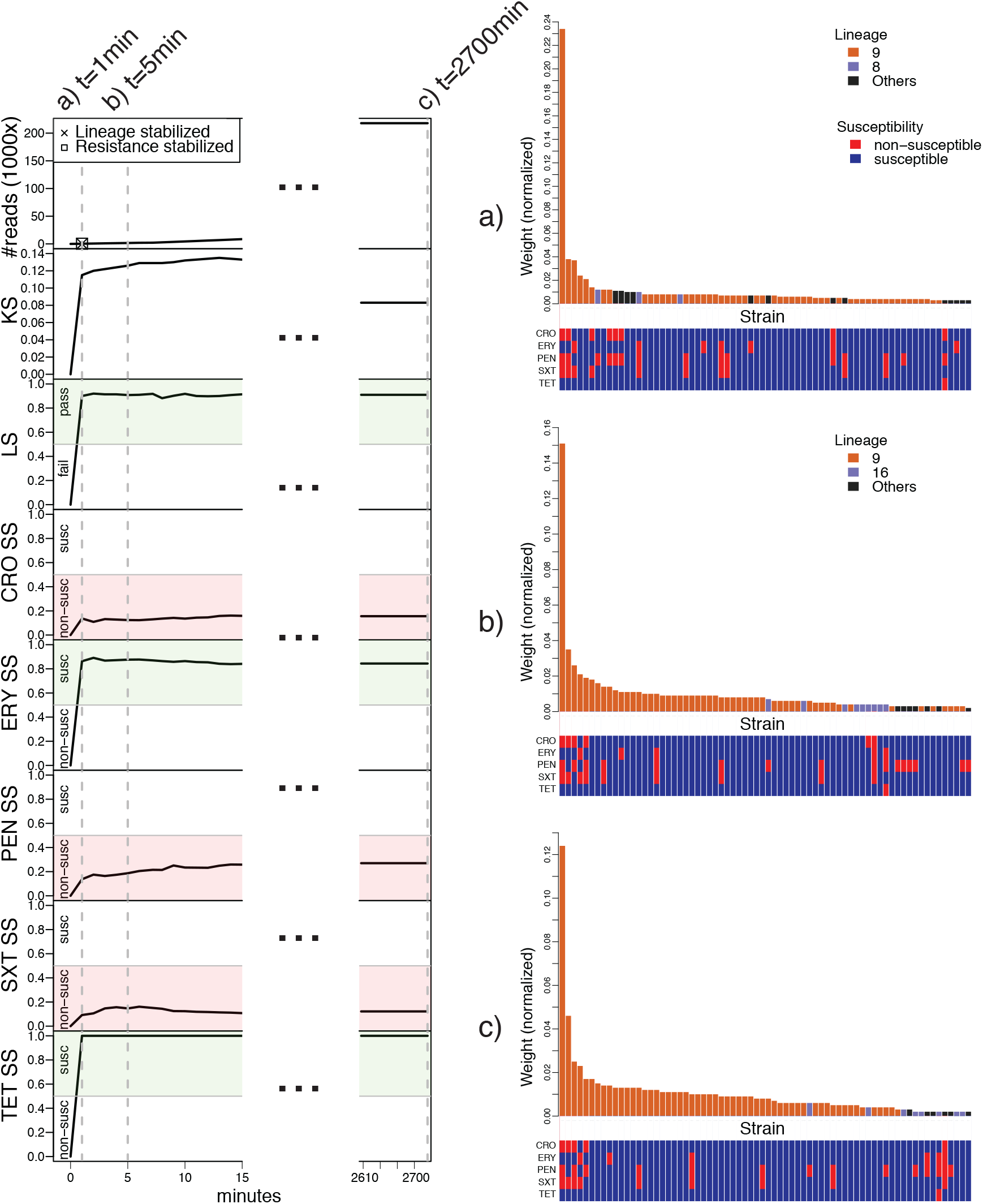
RASE obtains stable predictions of antibiotic resistance or susceptibility and lineage within minutes for an isolate of a pneumococcal 23F clone (SP06). **Left:** Number of reads, lineage score (LS), *k*-mer score (KS), and susceptibility scores (SS) for individual antibiotics as a function of time from the start of sequencing. In the top left plot, the times of stabilization are shown for the predicted lineage and susceptibility or resistance to all antibiotics. **Right:** a)-c) Similarity rank plots for selected time points (1 minute, 5 minutes, and the end of sequencing). The bars correspond to 70 best matching strains in the database and display the normalized weights, which serve as a proxy to inverted genetic distance. They are arranged by rank and colored according to the presence in the predicted, alternative or another lineage. The bottom panels display the resistance profiles of the strains.

### RASE databases for hundreds of S. pneumoniae and N. gonorrhoeae strains

We constructed RASE databases for *S. pneumoniae* and *N. gonorrhoeae* from the same data as described above (Methods). We assigned each pneumococcal and gonococcal strains to an antibiotic-specific resistance categories using the European Committee on Antimicrobial Susceptibility Testing (EUCAST) breakpoints^33^ and the CDC Gonococcal Isolate Surveillance Project (GISP) breakpoints^34^, respectively (Methods). Where MIC data were unavailable or insufficiently specific, we estimated the likely resistance phenotype using ancestral state reconstruction (Methods, **Supplementary Note 1**). To verify the results, we tested eight pneumococcal isolates for which resistance phenotypes were not originally available (Methods), and the measured MICs by microdilution matched the expected phenotypes (shown in bold in **Table 1**). We constructed the ProPhyle *k*-mer indexes with a *k*-mer length optimized to minimize prediction delays (*k*=18, Methods). The obtained pneumococcal and gonococcal RASE databases occupy 321 MB and 242 MB RAM (4.3× and 12× compression rate) and can be further compressed for transmission to 47 MB and 32 MB (29× and 90x compression rate), respectively (**Supplementary Figure 1**). This would allow RASE to be used on portable devices and its databases easily transmitted to the point of care over links with a limited bandwidth.

**Table 1:**
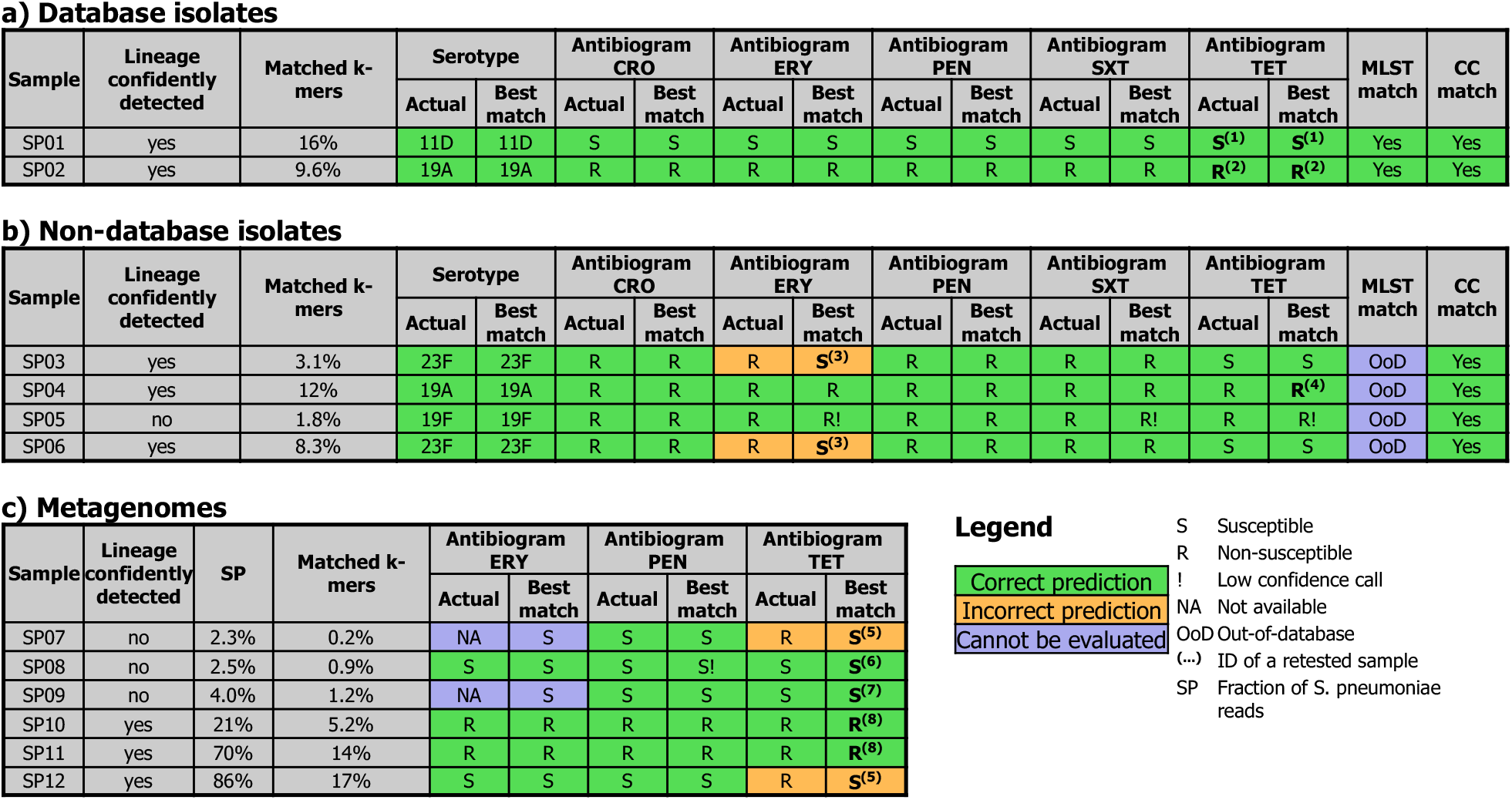
Predicted phenotypes of S. pneumoniae for a) database isolates, b) non-database isolates, and c) metagenomes. The table displays actual and predicted resistance phenotypes (S = susceptible, R = non-susceptible) for individual experiments, as well as information on match of the predicted MLST sequence type and clonal complex. Resistance categories in bold were inferred using ancestral reconstruction and were also confirmed using phenotypic testing (see Methods and Supplementary Table 3). Metagenomic samples are sorted by the estimated fraction of *S. pneumoniae* reads.

### RASE identifies strains in the database within minutes

We first examined two pneumococcal isolates that were used to build the RASE database (**Table 1a**, sens/spec 100%/100%, n=10) to test RASE can function in ideal circumstances. In the case of a fully susceptible isolate (SP01), the correct lineage and sequenced strain were identified within 1 minute and 7 minutes respectively. A multidrug-resistant isolate (SP02) was predicted even faster, with both lineage and the sequenced strain correctly detected and stabilized within 1 minute. To compare with gene-based approaches for detecting resistance^22^ we evaluated how long it took for resistance genes to be sequenced on the device, and observed that at least 25 minutes would be needed for single copies to be detected (**Supplementary Note 2**).

We then performed a similar evaluation with five gonococcal isolates from the database (**Table 2a**, sens/spec 57%/100%, n=20); however, here we selected more complicated cases. First, we tested a susceptible isolate (GC01), for which RASE identified the correct strain and antibiogram within 3 minutes of sequencing. We then sequenced an isolate with a novel and uncommon mechanism of cephalosporin resistance that has emerged recently (GC02)^35^. Under such circumstances, the resistant strain and its susceptible neighbors tend to be genetically very similar, which could confound our analysis. However, RASE was still able to identify the correct resistance phenotypes in 9 minutes, with the delay being due to difficulty distinguishing between the close relatives, reflected also by a susceptibility score in the low-confidence range (Methods). This was repeated in further experiments with the same isolate (GC03) which consistently reported low confidence in resistance phenotype (Methods), which is a feature of our approach intended to draw operators’ attention and indicate that further testing is necessary. In this experiment, RASE also resolved sample mislabeling (**Supplementary Note 3**). For a multidrug-resistant isolate (GC04) RASE predictions stabilized within 2 minutes but incorrectly predicted susceptibility to ceftriaxone. A subsequent analysis revealed that the ceftriaxone MIC of the sample was equal to the CDC GISP breakpoint (0.125 μg/mL), whereas the best match in the database had an MIC of 0.062 μg/mL, within a single doubling dilution. We further found that RASE performed well even with extremely poor data and low-quality reads (GC05, **Supplementary Note 4**). We also evaluated how genomic neighbor typing would perform if RASE used Kraken^31^ instead of ProPhyle^28^ (**Supplementary Note 5**).

**Table 2:**
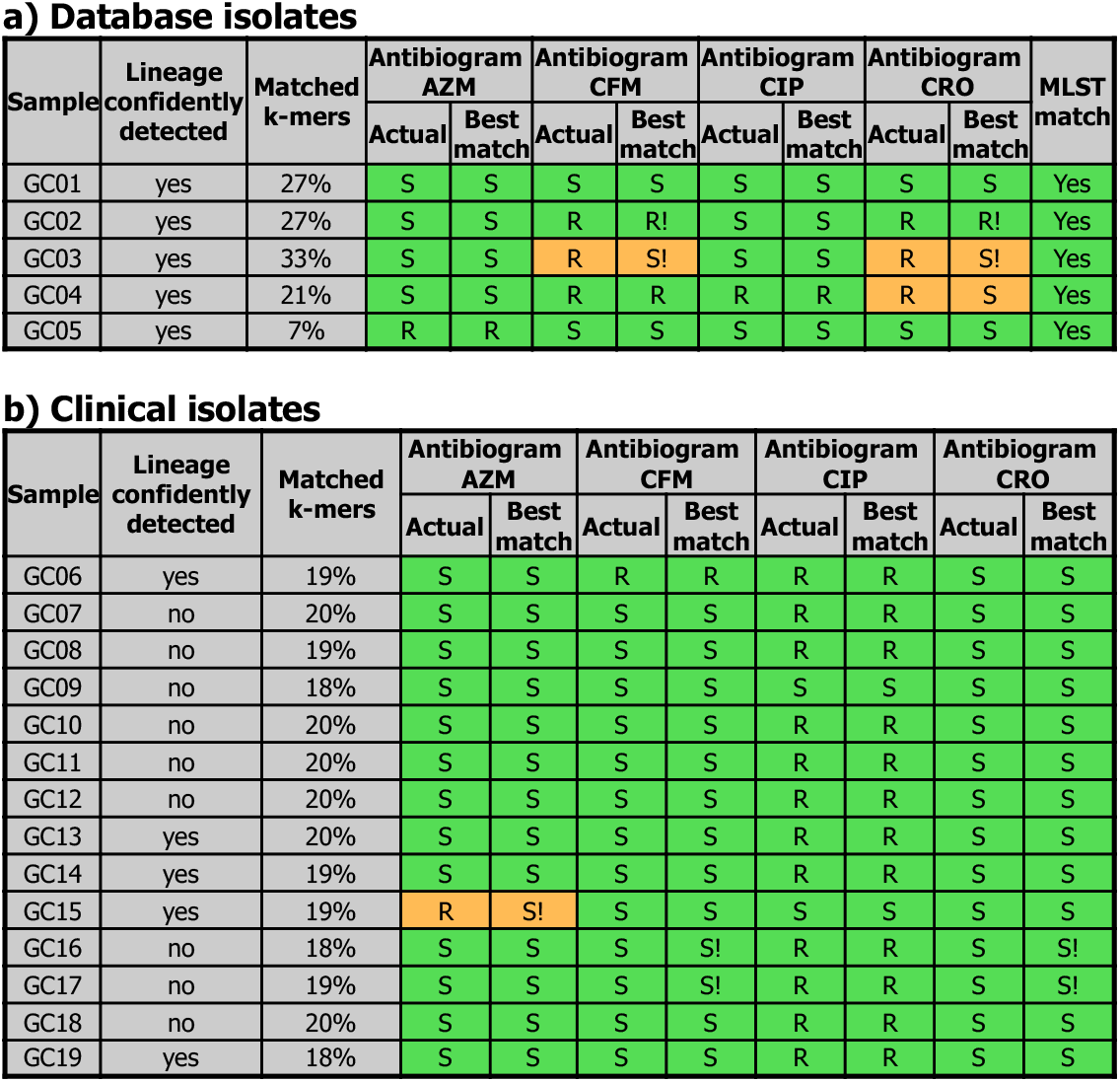
Predicted phenotypes of *N. gonorrhoeae* for a) database isolates and b) clinical isolates. The table is in the same format as Table 1.

### RASE identifies the closest relative of novel isolates

We next examined four novel pneumococcal isolates (**Table 1b**, sens/spec 89%/100%, n=20) for which the serotype and limited antibiogram and lineage data were known. We compared three characteristics of the sample to assess our performance: the serotype, the MLST sequence type, and the antibiograms (benzylpenicillin, ceftriaxone, trimethoprim-sulfamethoxazole, erythromycin, and tetracycline resistance according to the EUCAST breakpoints^33^).

In all cases, the closest relative was identified within 5 minutes, even if the correct MLST sequence type was absent from the RASE database (an example shown in **Figure 2**). The two samples from the 23F clone (SP03 and SP06) were correctly called as being closely related to the Tennessee 23F-4 clone identified by PMEN, a clone strongly associated with macrolide resistance^36^. Consistent with this, the two samples were indeed resistant to erythromycin. However, the Tennessee 23F-4 clone was absent from the Massachusetts sample, with the best match being a comparatively distantly related strain that was penicillin resistant, but erythromycin susceptible. This illustrates the importance of a relevant database.

We evaluated RASE with 14 clinical gonococcal isolates from the RaDAR-Go project^37^ (Switzerland, 2015–2016) (**Table 2b**, sens/spec 93%/100%, n=56). These isolates were previously sequenced using nanopore and have full antibiograms available^38^. The 55/56 correct calls indicate the strength of the genomic neighbor typing in a clinical setting. The only incorrect call (susceptibility to azithromycin in GC15) was marked as being low-confidence call on the basis of a poor susceptibility score. It should be noted that the ranges for what is considered low-confidence could vary among settings and pathogens but can be empirically determined and modified by users. In this case our results suggest that informative results can be obtained even using a database from one region (the US) to predict phenotype in another (Europe). However, this may not be the case for all pathogens.

### Phenotyping is still informative but lower quality on highly divergent lineages

As noted above, an important precondition of genomic neighbor typing is a comprehensive and relevant reference database. To evaluate RASE performance in a setting with an incomplete database, we used the gonococcal WHO 2016 reference strain collection^39^. This includes a global collection of 14 diverse isolates from Europe, Asia, North America, and Australia, collected over two decades and exhibiting phenotypes ranging from pan-susceptibility to multidrug resistance, and as such the GISP database is expected to be non-representative in this study. The WHO strains are available from the National Collection of Type Cultures, and were previously sequenced using nanopore^38^ and genetically and phenotypically characterized^39^. Surprisingly, RASE correctly identified all MLST sequence types represented in the database and in 7 cases it provided fully correct resistance phenotypes (**Supplementary Table 1**, sens/spec 67%/91%, n=56). In 6/7 cases where the complete resistance profile was not recovered, the closest relatives were identified correctly but were genetically divergent from the query isolates (**Supplementary Note 6**). In one case, the errors were due to a misidentification of the closest relatives by ProPhyle. Therefore, most prediction errors could be addressed with a more comprehensive database.

### RASE can identify resistance in pneumococcus from sputum metagenomic samples

Because bacterial culture and phenotyping via agar-dilution, Etest, or disk diffusion introduces significant delays in resistance profiling, direct metagenomic sequencing of clinical samples would be preferable for point-of-care use. We therefore analyzed metagenomic nanopore data from sputum samples obtained from patients suffering from lower respiratory tract infections^40^ (UK, 2017), selecting 6 samples from the study that were already known to contain *S. pneumoniae* (**Table 1c**, sens/spec 75%/100%, n=16).

One sample (SP10) contained DNA from multiple bacterial species. However, within 5 minutes sequence was identified belonging to the Swedish 15A-25 clone (ST63) which is also known to be associated with resistance phenotypes including macrolides and tetracyclines^41^. This sample was confirmed to be resistant to erythromycin, as well as clindamycin, tetracycline and oxacillin according to the EUCAST breakpoints^33^. The original report of the Swedish 15A-25 clone did not report resistance to penicillin antibiotics^41^, which has subsequently emerged in this lineage. However, our database correctly identified the risk of penicillin resistance in this sample. The metagenomes SP11 and SP12 contain an estimated >20% reads that matched to *S. pneumoniae*, and their serotypes were identified to be 15A and 3, respectively. The susceptibility scores of the best matches were fully consistent with the resistance profiles found in the samples, with the exception of tetracycline resistance in SP12 due to an incomplete database (**Supplementary Note 7**). The last remaining samples, SP07–SP09, contained less than 5% unambiguously pneumococcal reads. Despite the low proportions, all predicted phenotypes were concordant with phenotypic tests, with the exception of SP07, which matched the same strain as SP12 (discussed above).

## Discussion

This paper presents a method that we term genomic neighbor typing to pinpoint the closest relatives of a query genome within a suitable database and then to infer the phenotypic properties of the query strain on the basis of the reported properties of its relatives. At present, the precise lineage of a bacterial pathogen is often determined after most important clinical decisions have been made. However, incorporating genomic neighbor typing at an earlier stage offers a way of leveraging bacterial population structure to gain information on resistance and susceptibility, and inform antimicrobial therapy. The results from the metagenomic samples suggest that it is possible to apply this approach directly to clinical samples, and the success with both *S. pneumoniae* and *N. gonorrhoeae* indicates that it may have wide application.

The two pathogens studied here present contrasting features; the gonococcus is Gram-negative, harbors plasmids, and has a strikingly uniform core genome, while the pneumococcus is Gram-positive, does not contain plasmids and is diverse in both its core and accessory genome. Both exhibit high rates of homologous recombination, which is expected to both spread chromosomally encoded resistance elements and to scramble the phylogenetic signal that we use to identify the lineages. Despite these differences and the large degree of recombination, our approach performs well with both pathogens, with some differences that indicate opportunities and limitations for the application.

The initial identification of the closest relative is consistently more robust in the pneumococcus than the gonococcus, as a result of the former having more *k*-mers that are specific to an individual lineage, reflecting greater sequence diversity. As a consequence of the much lower diversity in gonococcus, when multiple closely related genomes are present in the database, RASE fluctuates between them, even though it correctly identifies the region of the phylogeny. If these genomes vary in their resistance profile, this is properly reflected in an uncertain susceptibility score indicating that caution and further investigation are merited (e.g., GC03).

As in all inference, the principle limitation of genomic neighbor typing is the representativeness of the database. While we have made use of relatively small samples from limited geographic areas to demonstrate proof of principle, in practice there are multiple examples of large genome databases generated by public health agencies, which could be combined with metadata on resistance for genomic neighbor typing. Such databases could, if necessary, be supplemented with local sampling. The relevant question for our approach therefore becomes whether the database contains a sufficiently high proportion of strains that will be encountered in the clinic and whether the resistance data are correct. Further work is required to determine the optimal structure and contents of databases for each application, but we emphasize the range of pathogens which appear to show promise for this approach. These include *E. coli*, in which data on MLST type supplemented with epidemiologic information can consistently produce AUCs in excess of 0.90 for multiple antibiotics^42^, suggesting great potential for neighbor typing to offer excellent resolution superior to MLST. However, genomic neighbor typing may be less suitable in the case where there is little genomic variation (e.g., *Mycobacterium tuberculosis*) or when resistance emerges rapidly on independent and diverse genomic backgrounds (e.g., *Pseudomonas aeruginosa* or resistance elements on highly promiscuous plasmids).

In the case where the infectious agent is unknown this problem is significantly more challenging. *K*-mers from one pathogen can match others and produce false predictions, and so choice of the correct database for prediction is key. Doing this will likely require a two-step solution in which the reads are first passed through a metagenomic classifier such as Centrifuge^43^ or MetaMaps^44^, which would be used to select the correct RASE database on which to make a resistance call.

Another limitation is the time required for sample preparation, which currently includes human DNA depletion, DNA isolation, and library preparation, taking a total of 4 hours. This is a rapidly evolving area of technology and automated rapid library preparation kits are already in development^45^. Further advances in this space, in particular for the preparation of metagenomic samples, will be required to bring the method closer to the bedside.

We have demonstrated that effectively predicting resistance and susceptibility from sequencing data does not require knowledge of *causal* resistance determinants. In fact, neighbor typing only requires that the phenotype be sufficiently strongly associated with the population structure to make reliable predictions.

A key advantage of this approach is that it requires very little genomic data, thus it is not limited by high error rates or low coverage. In particular, it is not attempting to define the exact genome sequence of the sample being tested, but merely which lineage it comes from. As a result, even when a small fraction of *k*-mers in the read are informative in matching to the RASE database, this is sufficient to call the lineage. This has the benefit of being faster than gene detection by virtue of the informative *k*-mers being distributed throughout the genome, and so more likely to appear in the first few reads sequenced by the nanopore. Therefore, the approach we present here can be seen as an application of compressed sensing: by measuring a sparse signal distributed broadly across our data we can identify it with comparatively few error-tolerant measurements.

Genomic neighbor typing can also be used to detect other phenotypes that are sufficiently tightly linked to a phylogeny, such as virulence. Further applications may include rapid outbreak investigations, as the closely related isolates involved in the outbreak would all be predicted to match to the same strain in the RASE database. The approach also lends itself to enhanced surveillance, including in the field; the 2014–2016 Ebola outbreak in West Africa, for example, saw MinION devices used in remote locations without advanced healthcare facilities^2^. Finally, at present empiric treatment decisions are made within successive ‘windows’^46^, in which increasing information becomes available, from initial Gram stain to full phenotypic characterization. The information from genomic neighbor typing is a natural complement to this process with the potential to improve therapy long before it would become clinically apparent that the patient is not responding or before phenotypic susceptibility data were available. The combination of high-quality RASE databases with genomic neighbor typing offers an alternative forward-looking model for diagnostics and surveillance, with wide applications for the improved clinical management of infectious disease.

## Methods

### Overview

RASE uses rapid approximate *k*-mer-based matching of long sequencing reads against a database of strains to predict resistance via neighbor typing. The database contains a highly compressed exact *k*-mer index, a representation of the tree population structure, and metadata such as lineage, resistance profiles, MLST sequence type and serotype. The RASE prediction pipeline iterates over reads from the nanopore sequencer and provides real-time predictions of lineage and resistance or susceptibility (**Figure 1**).

### Resistance profiles

For all antibiotics, RASE associates individual strains with a resistance category, ‘susceptible’ (S) or ‘non-susceptible’ (R). First, intervals of possible MIC values are extracted using regular expressions from the available textual antibiograms. For instance, ‘>=4’, ‘2’, and ‘NA’ would be translated to the intervals [4,+∞), [2,2], and [0,+∞), respectively. Then the acquired intervals are compared to the antibiotic-specific breakpoints (see below; **Supplementary Figures 3 and 4**). If a given breakpoint is above or below the interval, susceptibility or non-susceptibility is reported, respectively. However, no category can be assigned at this step if the breakpoint lies within the extracted interval, an antibiogram is entirely missing, it is insufficiently specific, or its parsing failed. Finally, missing categories are inferred using ancestral state reconstruction on the associated phylogenetic tree while maximizing parsimony (i.e., minimizing the number of nodes switching its resistance category; **Supplementary Figures 5 and 6**). When the solution for a node is not unique, non-susceptibility is assigned.

### Genomic neighbor typing

All reference strains in the database are associated with similarity weights that are set to zero at the start of the run. Each time a new read is read from the stream, *k*-mer-based matching is applied to identify the strains with the maximum number of matching *k*-mers (see below). Such strains are read’s nearest neighbors in the database according to the 1/(‘number of matched k-mers’) pseudodistance.

The weights of the nearest neighbors are then increased according to the ‘information content’ of the read, calculated as the number of matched *k*-mers divided by the number of nearest neighbors. Reads that do not match (i.e., 0 matching *k*-mers in the database) are not used in subsequent analysis. The computed matches are also used for updating the *k*-mer score (KS), which is the proportion of matched *k*-mers in all reads. KS helps to assess whether a sample is truly matching the database and predicting resistance for the database species makes sense.

The obtained weights serve as a proxy to inverted genetic distance and are used as a basis for the subsequent predictions of the lineage, and antibiotic resistance and susceptibility.

### Predicting lineage

A lineage is predicted as the lineage of the best matching reference strain, i.e., the one with the largest weight. The quality of lineage prediction is further quantified using a lineage score (LS), calculated as LS=2f/(f+t)-1, where f and t denote the weights of the best matches in the first (‘predicted’) and in the second best (‘alternative’) lineage, respectively. The values of LS can range from 0.0 to 1.0 with the following special cases: LS=1.0 means that all reads were perfectly matching the predicted lineage, whereas LS=0.0 means that the predicted and alternative lineages were matched equally well.

LS is used to measure how well a sample matching the identified lineage. If LS is higher than a specified threshold (0.6 in default settings), the call is considered successful. If the score is lower than this, the sample cannot be securely assigned to a lineage, and this should draw operators’ attention. Note that custom RASE databases may require a re-calibration of the threshold.

### Predicting resistance and susceptibility

Resistance or susceptibility are predicted for individual antibiotics independently, based the weights of the strains that belong to the predicted lineage. These are used to calculate a susceptibility score, which is further interpreted by comparing to pre-defined thresholds.

The susceptibility score is calculated as SS=s/(s+r), where s and r denote the weights of the best matching susceptible and best matching non-susceptible strains within the lineage. The values of SS can range from 0.0 to 1.0 with the following special cases: SS=0.0 and SS=1.0 mean that all reads match only resistant or susceptible strains in the lineage, respectively. In practice, this happens only if the lineage is entirely associated with resistance or susceptibility. SS=0.5 means that the best matching resistant and susceptible strains are matched equally well. As follows from the score definition, if SS is greater than 0.5, then the best matching strain is susceptible, otherwise it is non-susceptible.

SS is used for predicting resistance or susceptibility as well as for evaluating the prediction’s confidence. If SS is greater than 0.5, susceptibility to the antibiotic is reported, non-susceptibility otherwise. Hence resistance is predicted as the resistance of the best match. However, when SS is within the [0.4, 0.6] range, it is considered a low-confidence call, and as such it should draw operators’ attention; this usually indicates that resistance or susceptibility emerged recently in the evolutionary history and genomic neighbor typing may not be able to confidently distinguish between these similar, but phenotypically distinct, strains. Note that the thresholds above might require a further re-calibration, based on the specific database, antibiotics, and application of RASE.

### S. pneumoniae *RASE database*

The *S. pneumoniae* RASE database was constructed with the EUCAST breakpoints^33^ ([mg/L]): ceftriaxone (CRO): 0.25, erythromycin (ERY): 0.25, benzylpenicillin (PEN): 0.06, trimethoprim-sulfamethoxazole (SXT): 1.00, and tetracycline (TET): 1.00. While we have used the above values in the present work, others may be readily defined and the database rapidly updated. This is especially useful in the case where breakpoints may vary depending on the site of infection (as is the case with pneumococcal meningitis and otitis media, where lower MICs are considered to be resistant^33^).

The draft assemblies were downloaded from the SRA FTP server using the accession codes provided in Table 1 in ref^24^. The phylogenetic tree was downloaded from DataDryad (accession: ‘10.5061/dryad.t55gq’). The pneumococcal ProPhyle index was constructed with the *k*-mer size *k*=18.

The obtained *S. pneumoniae* RASE database including the code and source data is available from https://github.com/c2-d2/rase-db-spneumoniae-sparc.

### N. gonorrhoeae *RASE database*

The *N. gonorrhoeae* RASE database was constructed with the CDC GISP breakpoints^34^ ([mg/L]): azithromycin (AZM): 2.0, cefixime (CFM): 0.25, ciprofloxacin (CIP): 1.0, and ceftriaxone (CRO): 0.125. Before applying the breakpoints, azithromycin MICs for strains collected before 2005 were doubled in order to correct for the known inconsistencies of the phenotyping protocol due to a change in formulation of the commercial media^47^.

The draft assemblies and the phylogenetic tree were downloaded from Zenodo (accession: ‘10.5281/zenodo.2618836’). Three prevalent types of plasmids^48^ were downloaded from GenBank, localized in the GISP database using BLAST^49^, and removed from the dataset: the cryptic plasmid (‘pJD1’, GenBank accession ‘NC_001377.1’), the beta-lactamase plasmid (‘pJD4’, GenBank accession ‘NC_002098.1’), and the conjugative plasmid (‘pEP5289’, GenBank accession ‘GU479466.1’). The gonococcal ProPhyle index was constructed with the *k*-mer size *k*=18.

The obtained *N. gonorrhoeae* RASE database including the code and source data is available from https://github.com/c2-d2/rase-db-ngonorrhoeae-gisp.

### K-mer-based matching

Reads were matched against the RASE databases using the ProPhyle classifier^27, 28^ (commit b55e026) and its ProPhex component^50, 51^. ProPhyle index stores *k*-mers of all strains in a highly compressed form, reducing the required memory footprint. In the database construction phase, the strains’ *k*-mers are first propagated along the phylogenetic tree and then greedily assembled to contigs. The obtained contigs are then placed into a single text file, for which a BWT index is constructed^30^.

In the course of sequencing, each read is decomposed into overlapping *k*-mers. The *k*-mers are then searched in the BWT index by ProPhex using BWT search using a sliding window^50^. For every *k*-mer, the obtained matches are translated back on the tree. This provides a list of nodes whose descending leaves are the strains containing that *k*-mer. Finally, strains with maximum number of matched *k*-mers are identified for each read, and reported in the SAM/BAM format^52^.

### Optimizing k-mer length

The *k*-mer length is the main parameter of the classification. First, the subword complexity function^53^ of pneumococcus was calculated using JellyFish^54^ (version 2.2.10) (**Supplementary Figure 7**). Then, based on the characteristics of the function and the *k*-mer range supported by ProPhyle, the possible range of *k* was determined as in [17, 32]. For these *k*-mer lengths, RASE indexes were constructed and their performance evaluated using the RASE prediction pipeline and selected experiments. While RASE showed robustness to *k*-mer length in terms of final predictions, prediction delays differed (**Supplementary Figure 8**). Based on the obtained timing data, we set *k* to 18.

### Comparison to Kraken

For each RASE database, a fake NCBI taxonomy was generated from the database tree. Then a library was built using Kraken^31^ (v1.1.1, with default parameters) from the same FASTA files as used for building the RASE database. Finally, Kraken databases were constructed for both *k*=18 and *k*=31.

The obtained Kraken databases were used to classify reads from individual experiments. The obtained Kraken assignment were subsequently converted using an ad-hoc Python script to RASE-BAM (a subset of the BAM format^52^ used by RASE). Finally, RASE prediction was applied on the BAM files, with the use of the RASE database metadata, and the results compared with the results of the standard RASE with ProPhyle.

### Measuring time

To determine how RASE works with nanopore data generated in real time, the timestamps of individual reads extracted were using regular expressions from the read names. These were then used for sorting the base-called nanopore reads by time. When the RASE pipeline was applied, the timestamps were used for expressing the predictions as a function of time. The times of ProPhyle assignments were also compared to the original timestamps to ensure that the prediction pipeline was not slower than sequencing.

When timestamps of sequencing reads were not available (i.e., the gonococcal WHO and clinical samples), RASE estimated the progress in time from the number of processed base pairs. This was done by dividing the cumulative base-pair count by the typical nanopore flow, which we had previously estimated from SP01 as 1.43Mbps per second. However, such an estimated progress is indicative only, as it does not follow the true order of reads in the course of sequencing. As the nanopore signal quality tends to decrease over time (see the decrease of KS in **Figure 2** after t=15mins), the randomized read order provides results of lower quality than true real-time sequencing.

### Lower time estimates on resistance gene detection

A complete genome of the multidrug-resistant SP02 isolate was assembled from the nanopore reads using the CANU^55^ (version 1.5, with default parameters). Prior to the assembly step, reads were filtered using SAMsift^56^ based on the matching quality with the pneumococcal RASE database: only reads at least 1000bp long with at least 10% 18-mers shared with some of the reference draft assemblies were used. The obtained assembly was further corrected by Pilon^57^ (version 1.2, default parameters) using Illumina reads from the same isolate (taxid ‘1QJAP’ in the SPARC dataset^24^) mapped to the nanopore assembly using BWA-MEM^58^ (version 0.7.17, with the default parameters) and sorted using SAMtools^52^.

The obtained assembly was searched for resistance-causing genes using the online CARD tool^8^ (as of 2018/08/01). All of the original nanopore reads were then mapped using Minimap2^59^ (version 2.11, with ‘-x map-ont’) to the corrected assembly and resistance genes in the reads identified using BEDtools–intersect^60^ (version 2.27.1, with ‘-F 95’). Timestamps of the resistance-informative reads were extracted and associated with the genes. Only reads longer than 2kbp were used in the analysis.

### Evaluation of the N. gonorrhoeae WHO samples

To evaluate the predictions of the WHO samples, we inferred a phylogenetic tree from a data set comprising both the GISP isolates and the WHO isolates. First, reads were downloaded for the GISP isolates (NCBI BioProject: ‘PRJEB2999’ and ‘PRJEB7904’) and for the WHO isolates F–P (NCBI BioProject: ‘PRJEB4024’). For the WHO isolates U–Z, read data were simulated from the finished de-novo assemblies (NCBI BioProject: ‘PRJEB14020’) using Art-Illumina^61^ (version 2.5.1). Reads were mapped to the NCCP11945 reference genome (GenBank accession: ‘CP001050.1’) using BWA-MEM^58^ (version 0.7.17) and deduplicated using Picard^62^ (version 2.8.0). Pilon^57^ (version 1.16, with ‘--mindepth 10 --minmq 20’) was used to call variants and further filtered to include only ‘pass’ sites and sites where the alternate allele was supported with AF > 0.9. Gubbins^63^ (version 2.3.4) with RAxML^64^ (version 8.2.10) were run on the aligned pseudogenomes to generate the final recombination-corrected phylogeny (**Supplementary File 1**).

The closest relatives identified by RASE were verified using the obtained tree. For every WHO isolate, the obtained RASE prediction was compared to the closest GISP isolate on the tree.

### Library preparation

For isolates SP01-SP06, cultures were grown in Todd–Hewitt medium with 0.5% yeast extract (THY; Becton Dickinson and Company, Sparks, MD) at 37°C in 5% CO2 for 24 hrs. High-molecular-weight (>1 μg) genomic DNA was extracted and purified from cultures using DNeasy Blood and Tissue kit (QIAGEN, Valencia CA). DNA concentration was measured using Qubit fluorometer (Invitrogen, Grand Island NY). Library preparation was performed using the Oxford Nanopore Technologies 1D ligation sequencing kit SQK LSK108.

For experiments SP07-SP12, library preparation was performed using the ONT Rapid Low-Input Barcoding kit SQK-RLB001, with saponin-based host DNA depletion used for reducing the proportion of human reads. More details can be found in the original manuscript^40^.

For isolates GC01-GC05, cultures were grown on Chocolate-Agar media i.e., Difco GC base media containing 1% IsoVitaleX (Becton Dickinson Co., Franklin Lakes, NJ) and 1% Remel Hemoglobin (Thermo Fisher Scientific, Carlsbad, CA) at 37°C in 5% CO2 for 20 hrs. For GC01-GC04 genomic DNA was extracted and purified from cultures using the PureLink Genomic DNA MiniKit (Thermo Fisher Scientific, Carlsbad, CA), and for GC05 DNA was extracted using the phenol-chloroform method^65^. Genomic DNA was extracted and purified from cultures using the PureLink Genomic DNA MiniKit (Thermo Fisher Scientific, Carlsbad, CA). DNA concentration was measured using the Qubit fluorometer (Invitrogen, Grand Island, NY). Library preparation was performed using the Oxford Nanopore Technologies 1D ligation sequencing kit SQK-LSK109.

### MinION sequencing

Sequencing was performed on the MinION MK1 device using R9.4/FLO-MIN106 flow cells, according to the manufacturer’s instructions. For experiments SP01-SP06, base-calling was performed using ONT Metrichor (versions 1.6.11 (SP01), 1.7.3 (SP02), 1.7.14 (SP03–SP06)) simultaneously with sequencing and all reads passing Metrichor quality check were used in the further analysis. For experiments SP07-SP12, the ONT MinKNOW software (versions 1.4-1.13.1) was used to collect raw sequencing data and ONT Albacore (versions 1.2.2-2.1.10) was used for local base-calling of the raw data after sequencing runs were completed. For experiments GC01– GC05, ONT MinKNOW software was used to collect raw sequencing data and ONT Albacore (version 2.3.4) was used for local base-calling.

### Testing resistance phenotype

Additional retesting of SPARC isolates was done using microdilution. Organism suspensions were prepared from overnight growth on blood agar plates to the density of a 0.5 McFarland standard. This organism suspension was then diluted to provide a final inoculum of 105 to 106 CFU/mL. Microdilution trays were prepared according to the NCCLS methodology with cation-adjusted Mueller-Hinton broth (Sigma-Aldrich) supplemented with 5% lysed horse blood (Hemostat Laboratories)^66, 67^. Penicillin (TRC Canada) and chloramphenicol (USB) concentrations ranged from 0.016 to 16 μg/mL. Erythromycin (Enzo Life Sciences), tetracycline (Sigma-Aldrich), and trimethoprim-sulfamethoxazole (MP Biomedicals) concentrations ranged from 0.0625 to 64 μg/mL. Ceftriaxone (Sigma-Aldrich) concentrations ranged from 0.007 to 8 μg/mL. The microdilution trays were incubated in ambient air at 35°C for 24 h. The MICs were then visually read and breakpoints applied. A list of individual microdilution measurements and the obtained resistance categories is provided in **Supplementary Table 2**.

Resistance of streptococcus in the metagenomic samples (SP07–SP12) was determined by agar diffusion using the EUCAST methodology and breakpoints^33^. First, the inoculated agar plates were incubated at 37 °C overnight and then examined for growth with the potential for reincubation up to 48 hours. Then, the samples were screened to oxacillin: if the zone diameter r was >20mm, the isolate was considered sensitive to benzylpenicillin, otherwise a full MIC measurement to benzylpenicillin was done. Finally, the isolate was screened for resistance to tetracycline (r≥25mm for sensitive, r<22mm for resistant) and erythromycin (r≥22mm for sensitive, r<19mm for resistant); when the isolate showed intermediate resistance, a full MIC measurement was done.

Results for all tested samples – isolates and metagenomes – are summarized in **Supplementary Table 3**.

## Data, implementation and availability

RASE was developed using Python, GNU Make, GNU Parallel^68^, Snakemake^69^, and the ETE 3^70^ and PySam^52^ libraries, and was based on ProPhyle (commit b55e026). Bioconda^71^ was used to ensure reproducibility of the software environments. All code, the generated databases and other supplementary materials are available under the MIT license from https://github.com/c2-d2/rase-supplement. The analyses in the paper were performed with the following versions of the RASE databases: “*N. gonorrhoeae* GISP USA v1.4” and “*S. pneumoniae* SPARC USA v1.3”. Sequencing data for all experiments can be downloaded from Zenodo (accession: ‘10.5281/zenodo.3346055’); for the metagenomic experiments, only the filtered datasets (i.e., after removing the remaining human reads *in silico*) were made publicly available.

## Acknowledgements

This work was supported by the Bill & Melinda Gates Foundation (GCGH GCE OPP1151010, KB and WPH), NIH – National Institute of Allergy and Infectious Diseases (R01 AI106786-05, KB), the Canadian Institutes of Health Research (MFE 152448, RSL), the Canadian Institutes for Health Research (a fellowship grant, DRM), and the David and Lucile Packard Foundation (MB). This paper presents independent research funded by the National Institute for Health Research (NIHR) under its Programme Grants for Applied Research Programme (Reference Number RP-PG-0514-20018, JOG), the UK Antimicrobial Resistance Cross Council Initiative (MR/N013956/1, JOG), Rosetrees Trust (A749, JOG), the University of East Anglia (JOG, TC), and Oxford Nanopore Technologies (JOG, TC). Portions of this research were conducted on the O2 and Odyssey high-performance compute clusters, supported by the Research Computing Groups at Harvard Medical School and at the Harvard Faculty of Arts and Sciences, respectively. The authors thank Joshua Metlay for providing the test isolates for experiments SP03–SP06, which were collected as part of a population-wide surveillance study done in the Philadelphia region, supported by NIH (R01 AI46645), and to Brian J Arnold, Taj Azarian and Cristina M Herren for useful comments in various stages of this project.

## Transparency declarations

JOG received financial support for attending ONT and other conferences and an honorarium for speaking at ONT headquarters. JOG received funding and consumable support from ONT for TC’s PhD studentship.

## Supplementary notes

**Supplementary Note 1.** Out of all 616 pneumococcal strains (**Supplementary Table 4a**), after the ancestral reconstruction step 485 were associated with susceptibility to ceftriaxone, 484 to erythromycin, 341 to benzylpenicillin, 480 to trimethoprim-sulfamethoxazole, and 551 to tetracycline (**Supplementary Table 5a**). In case of gonococcus, ancestral reconstruction was needed only for cefixime (62 records affected). Out of all 1102 gonococcal strains (**Supplementary Table 4b**), 808 were associated with susceptibility to azithromycin, 833 to cefixime, 508 to ciprofloxacin, and 1033 to ceftriaxone (**Supplementary Table 5b**). In our subsequent experiments, if original MIC data were not available for the best match in the RASE database, the relevant strain was tested to confirm resistance phenotype (Methods).

**Supplementary Note 2.** We evaluated how long it took for resistance genes to be reliably detected in nanopore reads. For SP02 we observed that at least 25 minutes were needed to detect resistance (i.e., to observe all resistance genes at least once), assuming that the genes in question can be unambiguously identified in nanopore data despite the high per-base error rate, and that the presence of the loci is directly linked to the resistance phenotype (**Supplementary Figure 2**). If this is not the case (for example if resistance is conferred by a single SNP, requiring coverage with multiple reads), further delays would be expected. Thus, genomic neighbor typing can offer a time advantage compared to methods based on identifying the presence of resistance genes even in a sample of DNA from a purified isolate as opposed to a metagenome, potentially allowing for more rapid changes to antimicrobial therapy.

**Supplementary Note 3.** We originally attempted to evaluate a multidrug-resistant isolate (GCGS0938 in the GISP collection); however, RASE placed it onto a distant part of the phylogeny and identified it as GCGS0324 or GCGS1095. A subsequent analysis revealed that the sample was mislabeled and that it was indeed GCGS1095, i.e., the same strain as in GC02, although from a different stock.

**Supplementary Note 4.** We evaluated how RASE performs in extremely unfavorable sequencing conditions; we sequenced an isolate (GC05) from the GISP collection with the use of an expired flow cell (purchased in October 2017, expired in December 2017, and the sequencing done in April 2018). In consequence, we obtained only 3.5 Mbps of low-quality reads (only 7% of matching *k*-mers compared to 20% obtained in the other isolates) (GC05 in **Table 2a**). An experiment with such a low yield would normally be discarded; despite that RASE provided correct and stabilized predictions (once the first long read was obtained from the sequencer at t=21mins).

**Supplementary Note 5.** We evaluated how genomic neighbor typing would perform if RASE used Kraken^31^ instead of ProPhyle^28^ for the read-to-strain comparison (the matching step in **Figure 1**). Both tools use *k*-mer-based matching to assign sequencing reads to a phylogenetic tree, but with several key differences. Whereas Kraken stores for each *k*-mer the lowest common ancestor (LCA) only, assigns reads to the LCA of the best hits and ignores low-complexity *k*-mers, ProPhyle indexes all *k*-mers using an exact index and can thus resolve ambiguities both on the level of individual *k*-mers and read assignments.

To compare both tools, we implemented a RASE wrapper for Kraken (Methods) and applied that to the same read and database data. We then compared the final inference results obtained with Kraken (with *k*=18 and *k*=31) with the results obtained from the standard RASE pipeline (**Supplementary File 2**).

For *S. pneumoniae* and *N. gonorrhoeae*, the number of inference errors increased more than 1.5x and 1.7x, respectively (in case of both *k*-mer sizes). In the case of *N. gonorrhoeae*, RASE-Kraken showed large systematic biases in neighbor typing, assigning 16 (*k*=18) and 18 (*k*=31) out of the gonococcal 33 samples to a single strain (GCGS1028), whereas RASE-ProPhyle identified this strain only once. While in the WHO dataset the numbers of RASE-ProPhyle and RASE-Kraken errors were comparable (10 vs. 12 and 11), in the RaDAR-Go dataset it increased from 1 to 8 and 10. Overall, the obtained results suggest that Kraken is less suited for the use in genomic neighbor typing than ProPhyle.

**Supplementary Note 6.** We analyzed the results of the WHO gonococcal samples (**Supplementary Table 1**). First, we evaluated the RASE ability to predict MLST sequence types. In all cases, either RASE predicted the correct sequence type (n=9), or the true sequence type was not present in the reference database (n=5). The latter was the case only in the samples F through P, which belonged to the initial 2008 WHO reference panel and were collected primarily in the late 1990s, with the majority of specimens isolated from the Eastern Hemisphere^72^. The GISP database, comprising strains collected in the US from 2000–2013, may not be representative then of the circulating lineages in those regions during that time span, which could result in both sequence type and antibiogram prediction errors. However, we observed perfect prediction of sequence types in the additional 2016 WHO reference strains comprising U through Z that were collected in 2007 and onwards^39^.

We next sought to evaluate the resistance predictions. In 7 cases (F, K, N, O, P, U, W), the antibiograms were identified fully correctly; in 4 (G, V, X, Z) and 3 cases (L, M, Y) one and two mistakes were made, respectively. To explain these discrepancies, we inferred a recombination-corrected phylogenetic tree comprising the GISP database isolates as well as the WHO samples (**Supplementary File 1**). With the exception of G and Y, the WHO isolates and their respective RASE-predicted best matches were the closest GISP isolates, indicative of accurate matching by RASE. While branch lengths of L, M and V on the tree reveal that the corresponding parts of the phylogeny are not well sampled in the database, the X, Y, and Z samples emerged from lineages that are well-represented but have acquired an atypically high level of cephalosporin resistance. Whereas X and Z acquired a novel resistance-conferring mosaic penA allele^73^, Y acquired a novel active site mutation in the context of a pre-existing mosaic penA allele^74^. While both of these adaptations resulted in high-level resistance, these mutations also appear to incur fitness costs in vitro and in the gonococcal mouse model^75^. In line with this, these strains have only been sporadically observed in genomic surveillance of clinical isolates. These results highlight how ancestral or emerging resistant lineages may not be well-captured by sequence-based methods including RASE and emphasize the value of continuous updating of the RASE database for public health.

**Supplementary Note 7.** Further analysis of the reads from SP12 using Krocus^76^ suggested that the pneumococcal DNA present was from the ST180 clonal complex, and matched specifically either to the sequence type ST180 or ST3798. This is consistent with identification as serotype 3, because this clonal complex contains the great majority of strains with this capsule type, which historically has not been associated with resistance^77^. However, improved sampling and study of this lineage has recently found highly divergent subclades that are associated with resistance. These lineages were previously rare, and thus were less likely to be included in our database, but now are increasing in frequency^78^. In this case, ST3798 is found to be in clade 1B, which is notable for exhibiting sporadic tetracycline resistance. Again, the failure to match to this is a result of the original database not containing a suitable example for comparison.

---

**Supplementary Figure 1:**
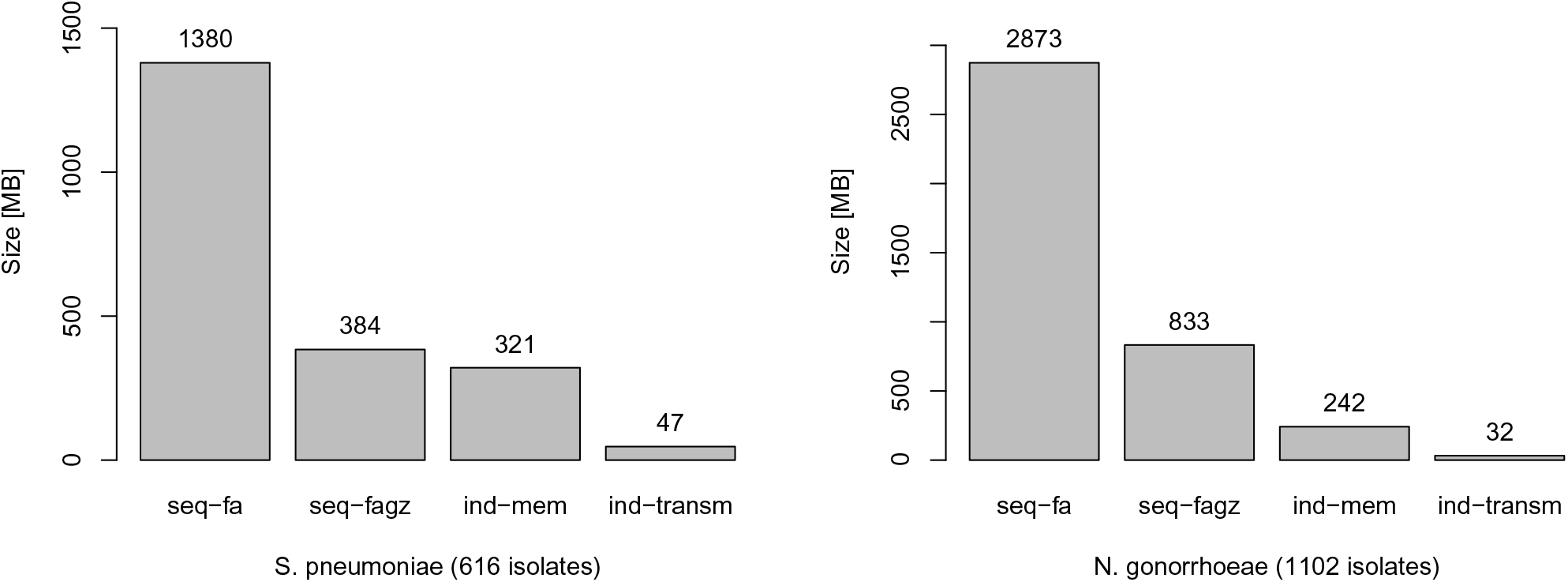
Size and memory footprint of the *S. pneumoniae* and *N. gonorrhoeae* RASE databases. The graph compares the size of the ProPhyle RASE index to the size of the original sequences: original draft assemblies (seq−fa), original draft assemblies compressed using gzip (seq−fagz), memory footprint of ProPhyle with the RASE index (ind−mem), and size of the ProPhyle RASE index compressed for transmission (ind−transm).

**Supplementary Figure 2:**
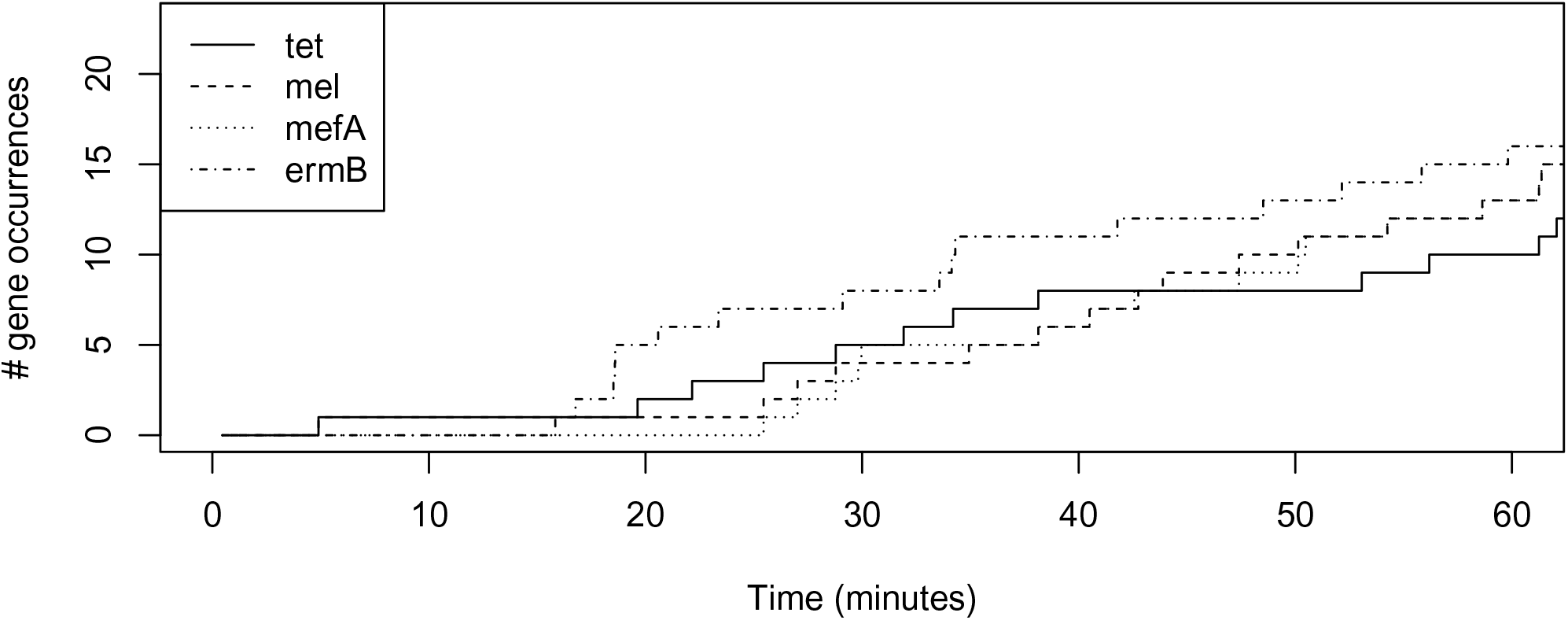
Timeline of resistance genes. Number of occurrences of individual resistance genes in reads of SP02, as a function of time for the first hour of nanopore sequencing.

**Supplementary Figure 3:**
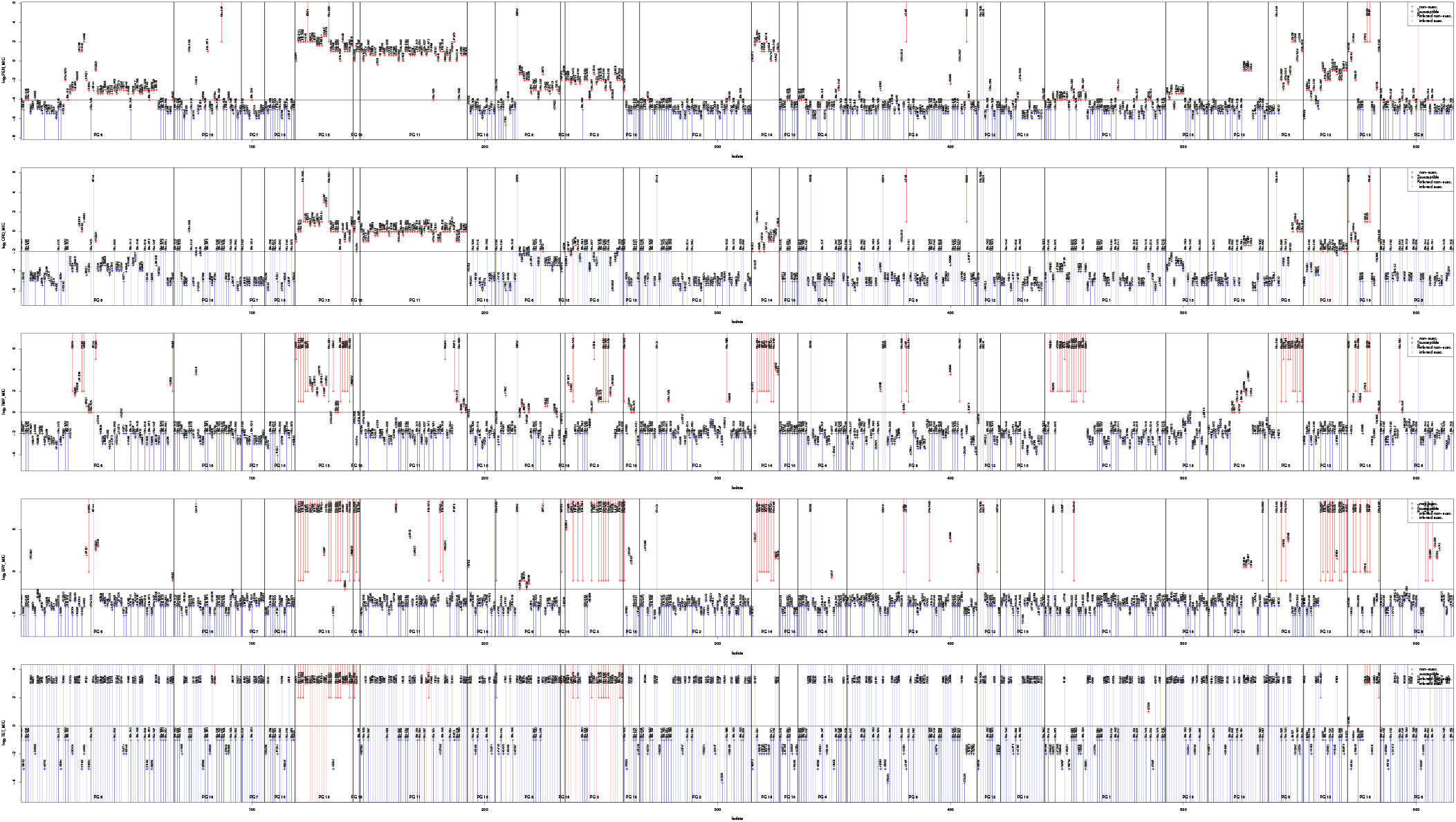
MIC intervals for individual strains in the *S. pneumoniae* RASE database. The plot illustrates MIC intervals and point values extracted from. Each panel corresponds to a single antibiotic, while vertical lines and points correspond to individual strains. Their colors correspond to the resistance category after applying a breakpoint (horizontal lines). When a resistance category could not be assigned directly (i.e., in case of an interval crossing the breakpoint line), then it was inferred using ancestral state reconstruction.

**Supplementary Figure 4:**
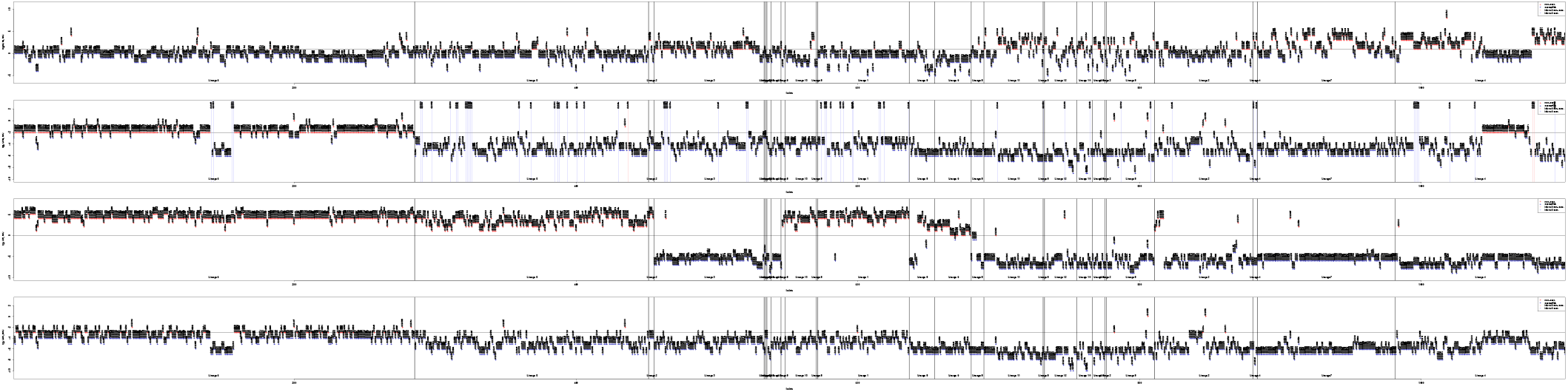
MIC intervals for individual strains in the *N. gonorrhoeae* RASE database. The figure is of the same format as Supplementary Figure 3.

**Supplementary Figure 5:**
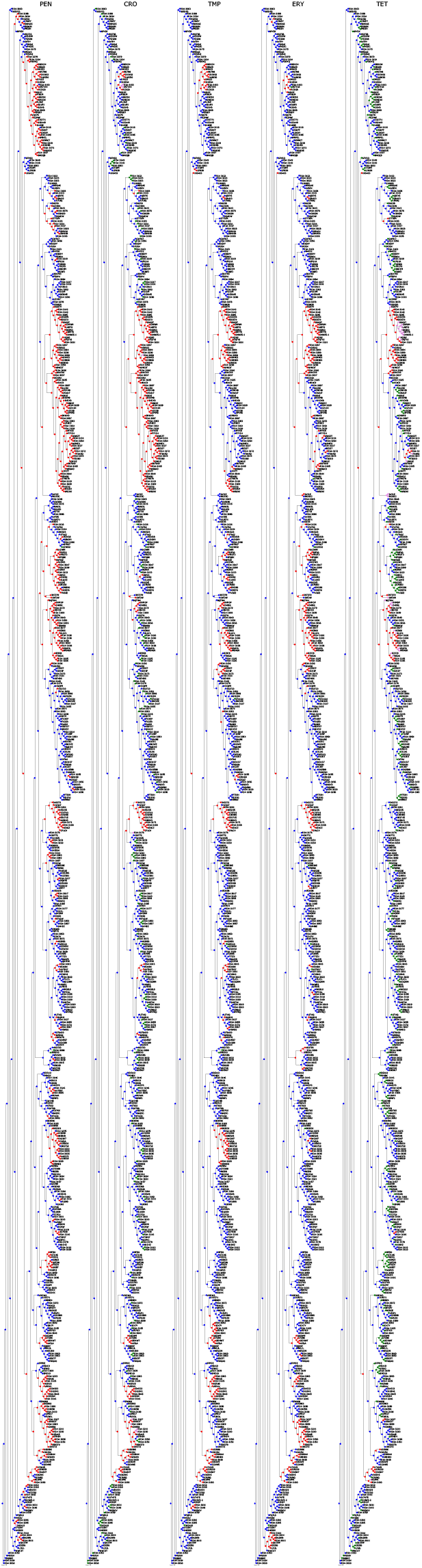
Ancestral state reconstruction of resistance categories in the *S. pneumoniae* RASE database. Each panel corresponds to a single antibiotic and displays the database phylogenetic tree, colored according to the reconstructed resistance categories for the antibiotic (blue, green, red, violet correspond to ‘susceptible’, ‘unknown – inferred susceptible’, ‘non-susceptible’, ‘unknown – inferred non-susceptible’, respectively).

**Supplementary Figure 6:**
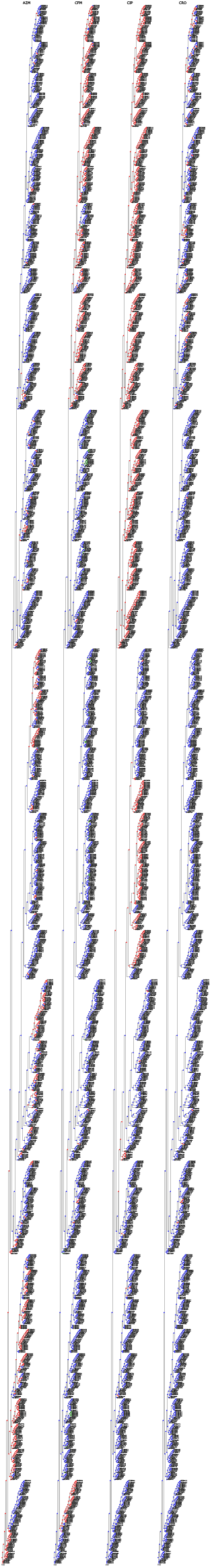
Ancestral state reconstruction of resistance categories in the *N. gonorrhoeae* RASE database. The figure is of the same format as Supplementary Figure 5.

**Supplementary Figure 7:**
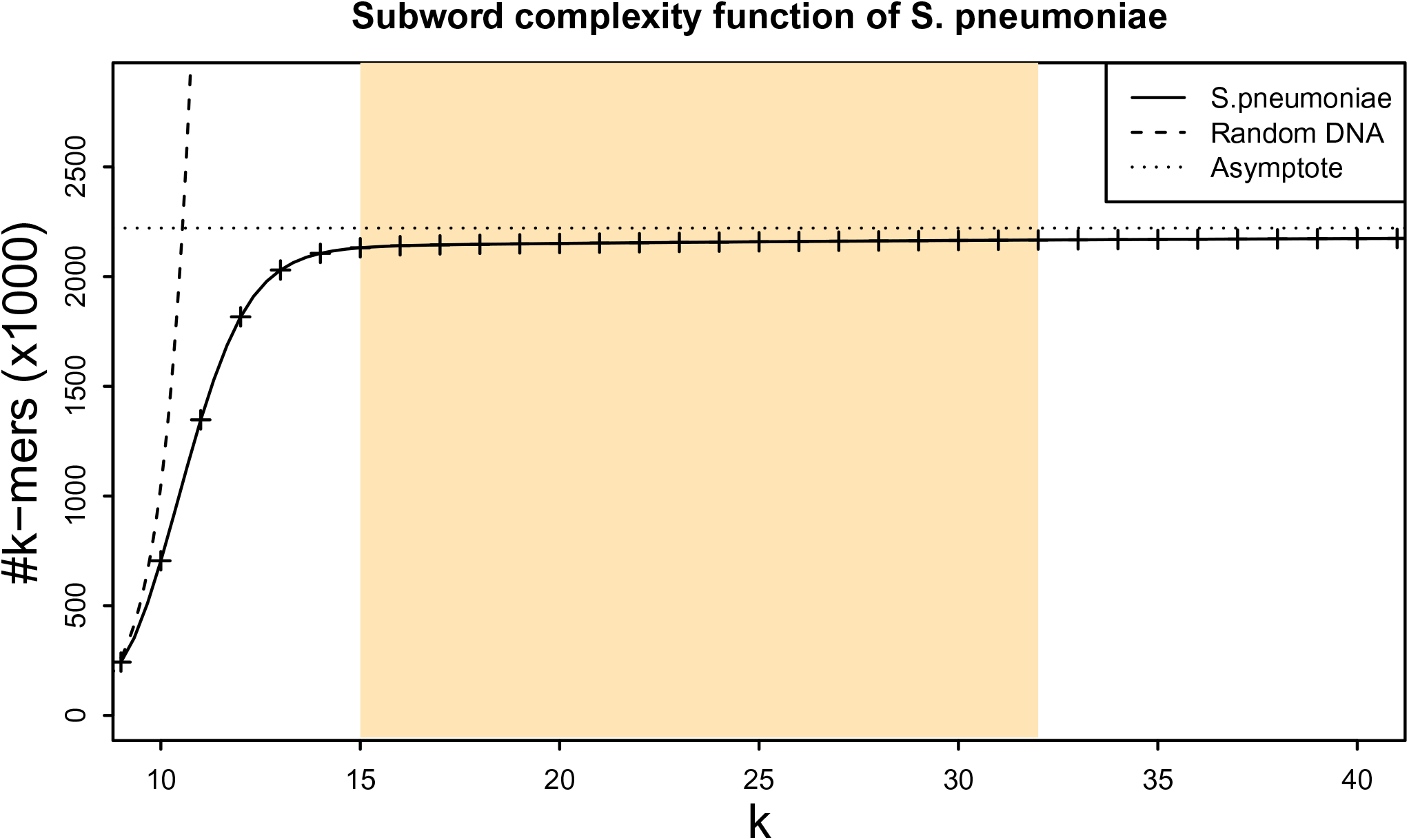
Subword complexity of pneumococcus. The plot depicts the number of canonical *k*-mers as a function of *k* for *S. pneumoniae* ATCC 700669 (GenBank accession: ‘NC_011900.1’) and for a random DNA text containing all possible *k*-mers. For *k <* 10, the pneumococcus *k*-mer composition is similar to the one of random text. For *k >* 14, the *k*-mer sets are almost saturated and the complexity grows very slowly. Since the genome length is finite and bacterial chromosomes are circular, the function attains its maximum at the genome size (2, 221, 315 in this case). The highlighted region corresponds to the range of values of *k*, which are suitable for use in RASE.

**Supplementary Figure 8:**
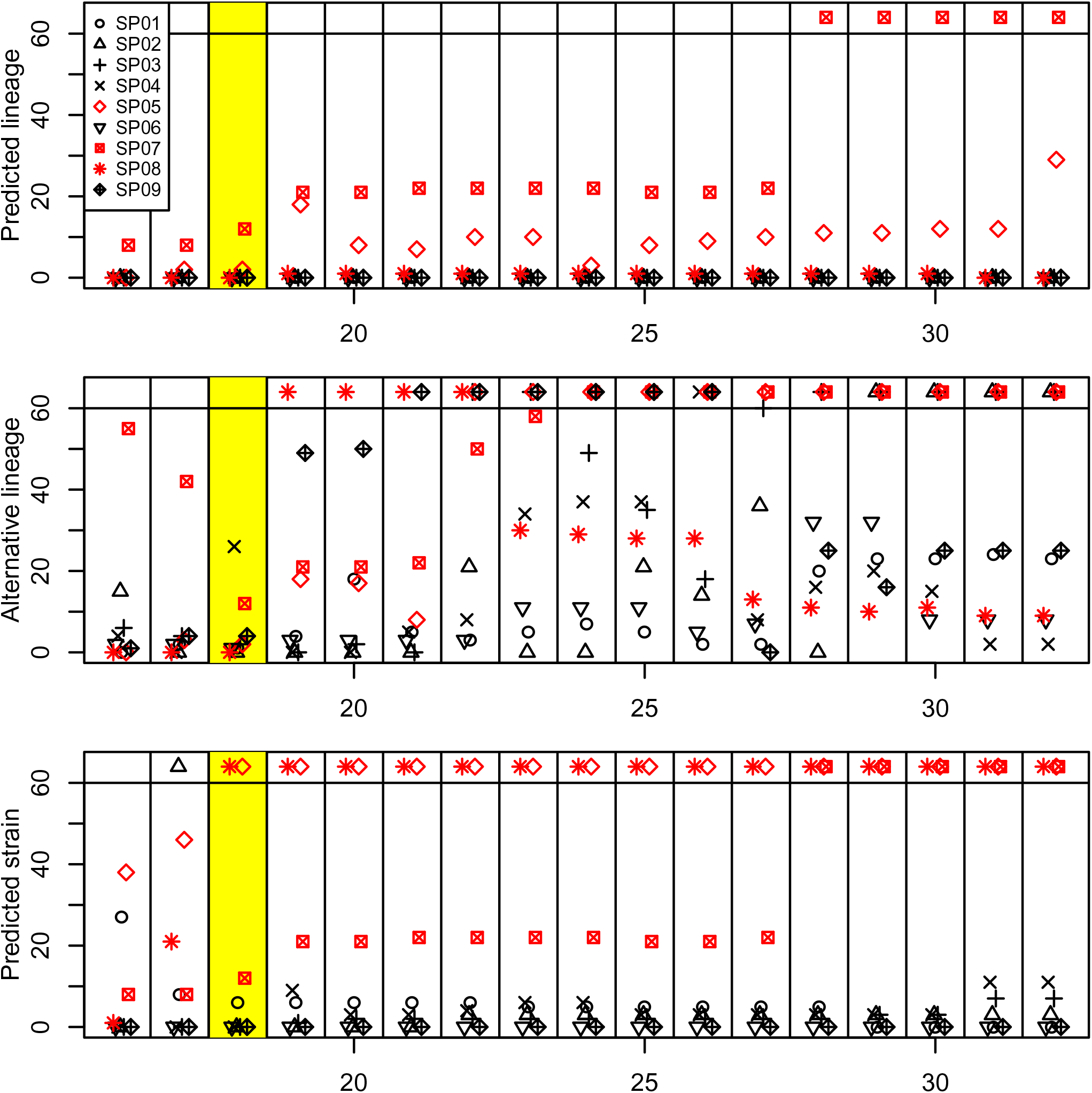
Delays in prediction based on the *k*-mer length. The plot displays delays in prediction as a function of the used *k*-mer length, for selected experiments and all possible *k*-mer lengths. Each horizontal panel displays times required for stabilization of one of the three predictions: the lineage, the alternative lineage, and the closest strain. Every column within a panel corresponds to a single *k*-mer length. When the required time exceeded 1 hour, the point is displayed at the top. Experiments where lineage could not be identified are plotted in red. The highlighted column corresponds to the *k*-mer length used for constructing the RASE databases in this paper.

**Supplementary File 1: Comprehensive phylogenetic tree for *N. gonorrhoeae***. A recombination-corrected tree in the Newick format comprising the GISP database isolates and the WHO samples.

**Supplementary File 2: Comparison of ProPhyle- and Kraken-powered genomic neighbor typing**. The spreadsheet shows the final resistance and susceptibility inference calls for the ProPhyle (k=18) and Kraken (k=18 and k=31) classifiers plugged into RASE; erroneous calls are highlighted in red.

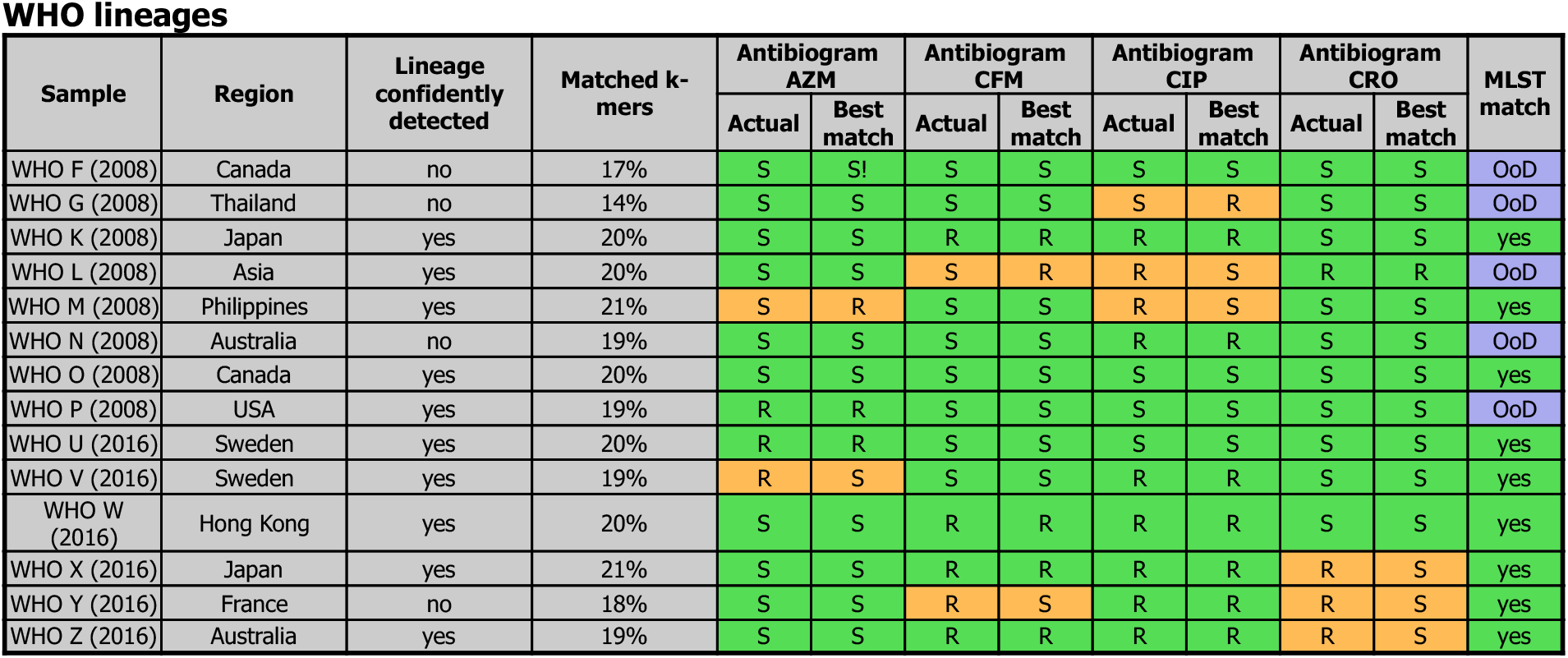

**Supplementary Table 1:** Predicted phenotypes of *N. gonorrhoeae* for the WHO lineages. The table is in the same format as Table 1.

**Supplementary Table 2: Additional MIC measurements for selected strains**. The table displays results from strain retesting. Each record contains date when the retesting was done, the antibiotic, the measured MIC, and the corresponding resistance category.

**Supplementary Table 3: Overview of performed resistance tests**. For all sequencing experiments, the table displays the best matching strains, their MICs, and all measurements of database MICs (the original reported values or categories inferred using ancestral state reconstruction when not available, retested values, and the resulting resistance categories).

**Supplementary Table 4: Metadata for all strains included in the a) *S. pneumoniae* and b) *N. gonorrhoeae* RASE database**. Each record contains the strain’s taxid, lineage, serotype (for S. pneumoniae only), MLST sequence type, order in the phylogenetic tree, and three fields related to resistance for every antibiotics: the ‘_mic’, ‘_int’, ‘_cat’ fields contain the original published MIC information (possibly corrected after retesting), the extracted MIC interval, and the resulting category after ancestral state reconstruction (S = susceptible, R = non-susceptible, s = unknown but reconstructed susceptible, r = unknown but reconstructed non-susceptible), respectively.

**Supplementary Table 5:** Prevalence of resistance phenotypes across lineages in the a) *S. pneumoniae* and b) *N. gonorrhoeae* RASE database. Statistics on prevalence of resistance phenotypes across lineages before and after the ancestral state reconstruction step.

**Supplementary Table 6: Sensitivity and specificity of resistance and susceptibility inference in all the datasets**. The table shows the number of true positive (TP), true negative (TN), false negative (FN), and false positive (FP) calls for resistance/susceptibility in individual datasets and the resulting sensitivity and specificity.

