## Supplementary Document 1 for "Rapid heuristic inference of antibiotic resistance and susceptibility by genomic neighbor typing"

### Quantification of the association of bacterial clones with antibiotic resistance

Supplementary Document 1 for 'Rapid heuristic inference of antibiotic resistance by genomic neighbor typing'

#### 1 Introduction

In this document, we show how to quantify the association of bacterial clones with antibiotic resistance, within and across species. For individual bacterial populations, we construct optimal predictors of resistance from lineages and calculate the associated Areas under the Curve (AUC).

We use this framework to show that for the pathogens *S. pneumoniae* and *N. gonorrhoeae* antibiotic resistance is highly associated with the population structure. This provides evidence of the suitability of genomic neighbor typing as a diagnostic method for these pathogens.

#### 2 Optimal lineage-to-resistance classifiers

We use the following assumptions and notation. Let us have a bacterial species and assume that it has $g$ equally probable lineages. Assume that we have $N$ isolates ($N \gg g$) for which we want determine resistance. Assume that every isolate belongs to exactly one lineage and we that for an isolate we can always correctly determine that lineage (denoted as $lineage(isolate)$). For every antibiotic, let us assume that each isolate is either resistant or susceptible, and the resistance phenotype within a lineage $i$ is iid with Bernoulli distribution, i.e., an isolate of the lineage $i$ is 'S' or 'R' with a probability $r_i$.

In order to mathematically quantify the association of resistance to the lineages, we construct memoryless probabilistic resistance classifiers of the form

$$C(lineage(sample)) \to \{\text{'S'}, \text{'R'}\}.$$

In other words, such a classifier predicts resistance only based on the knowledge of the lineage of the sample.

Such classifiers can be parametrized as $C_{R_1,\ldots,R_g}$, where $R_i \in [0,1]$ is the probability that resistance will be reported if the sample belongs to the lineage $i$. For instance, the classifier $C_{0,\ldots,0}$ always assigns 'S', $C_{\frac{1}{2},\ldots,\frac{1}{2}}$ assigns resistance like a fair coin, and $C_{1,\ldots,1}$ always assigns 'R'.

For every classifier $C_{R_1,\ldots,R_g}$, we can find an explicit formula for false positive rates (FPRs) and true positive rates (TPRs). Let us use the standard notation:

$$FPR = \frac{FP}{FP + TN} \qquad \text{and} \qquad TPR = \frac{TP}{TP + FN}$$

where

$$TP = \#\text{true positives, i.e., resistant isolates predicted as resistant}$$
$$FP = \#\text{false positives, i.e., susceptible isolates predicted as resistant}$$
$$FN = \#\text{false negatives, i.e., resistant isolates predicted as susceptible}$$
$$TN = \#\text{true negatives, i.e., susceptible isolates predicted as susceptible}$$

Given our assumptions, we can estimate the performance of a classifier $C_{R_1,\dots,R_g}$ as

$$\begin{array}{ll} TP \approx \frac{N}{g}\sum_{i=1}^{g} r_i R_i & FP \approx \frac{N}{g}\sum_{i=1}^{g}(1-r_i)R_i \\ FN \approx \frac{N}{g}\sum_{i=1}^{g} r_i(1-R_i) & TN \approx \frac{N}{g}\sum_{i=1}^{g}(1-r_i)(1-R_i) \end{array}$$

where $N$ denotes the number of samples tested, which provides the following estimates:

$$FPR \approx \frac{\sum_{i=1}^{g}(1-r_i)R_i}{\sum_{i=1}^{g}(1-r_i)} \qquad \text{and} \qquad TPR \approx \frac{\sum_{i=1}^{g} r_i R_i}{\sum_{i=1}^{g} r_i}$$

Using these formulae, we can map every classifiers $C_{R_1,\dots,R_g}$ to the ROC space by

$$f(R_1,\dots,R_g) \rightarrow \left(\frac{\sum_{i=1}^{g}(1-r_i)R_i}{\sum_{i=1}^{g}(1-r_i)}, \frac{\sum_{i=1}^{g} r_i R_i}{\sum_{i=1}^{g} r_i}\right)$$

It is easy to see that the map $f$ is linear.

Now let us observe that the set of all possible classifiers forms a cube $[0,1]^g$ in the parameter space $\mathbb{R}^g$. Since the map $f$ is linear and the cube is convex, its image in the ROC space must be convex too; an example is provided in Figure 1. Moreover, the image of the cube is equal to the complex hull of the images of individual cube vertices (red points in Figure 1).

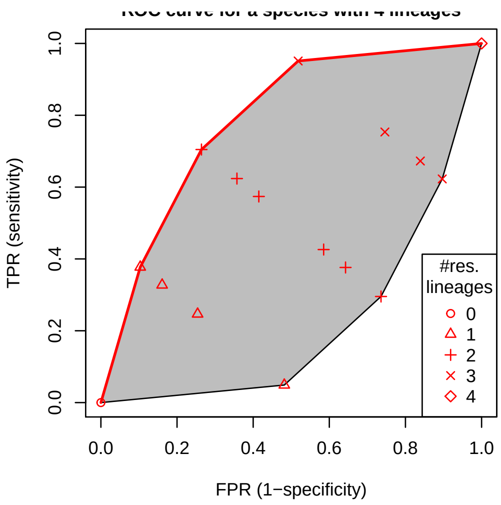


Figure 1: An illustration of the ROC diagram for a species with 4 lineages and a fixed antibiotic. The grey area corresponds to all the possible lineage-to-resistance classifiers $C_{R_1,R_2,R_3,R_4}$ while the red points are the classifiers with integer parameters, i.e., $R_i \in \{0,1\}$ for every $i \in \{1,\dots,g\}$. The optimal classifiers that maximize the AUC lie on the piece-wise linear red curve.

Our aim now is to find the optimal ROC curve maximizing the AUC (the red curve in Figure 1). Even though the curve corresponds to infinitely many classifiers, it is a piece-wise linear function fully defined by $g+1$ classifiers $C^{(0)},\dots,C^{(g)}$ lying on the line breaks. Enumerating this classifier sequence corresponds to putting lineages to the order in which they will be switched from susceptible to resistant (i.e., $R_i = 0 \rightarrow R_i = 1$). The first classifier $C^{(0)}$, $(0,0)$ in the ROC diagram, corresponds to all lineages being marked as susceptible, whereas the last classifier, $C^{(g)}$, $(1,1)$ in the diagram to all lineages being predicted marked as resistant.

The optimal ROC curve can be computed in multiple different ways. For instance, we can enumerate all cube vertices in the parameter space, map them to the ROC space, compute the convex hull, and extract its upper part. More efficiently, we can construct the classifier sequence directly in the ROC diagram by the following iterative process. We start at $(0,0)$, i.e., with all lineages marked as susceptible, and at every step we switch one susceptible lineage to resistant so that the corresponding step in the ROC graph has maximal possible slope, and we continue until all lineages are marked as resistant.

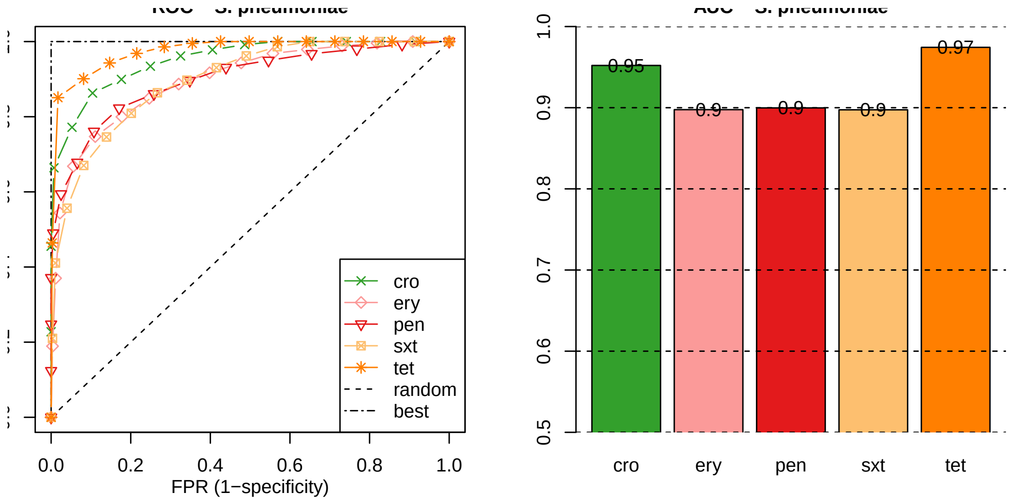


Figure 2: *S. pneumoniae* ROC curves and the corresponding AUCs for ceftriaxone (cro), erythromycin (ery), penicillin (pen), trimethoprim (sxt), and tetracycline (tet).

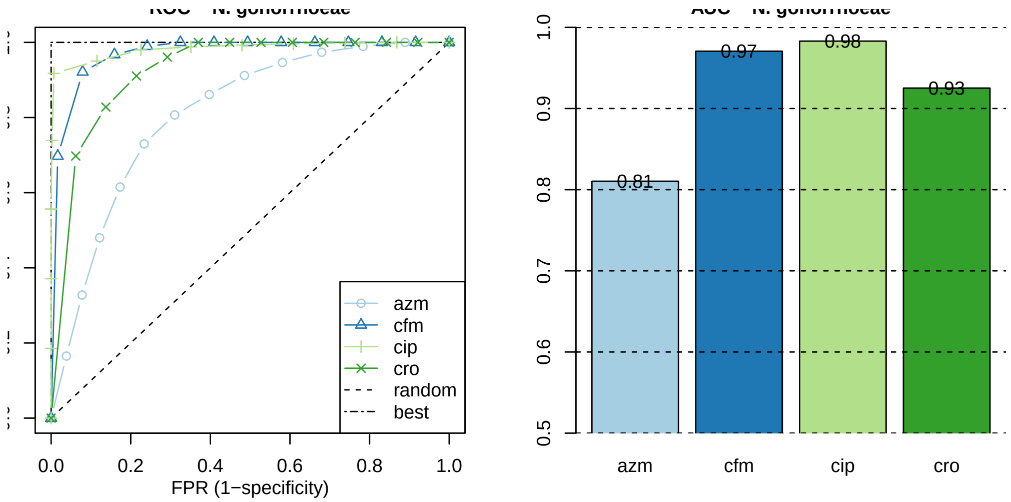


Figure 3: *N. gonorrhoeae* ROC curves and the corresponding AUCs for azithromycin (azm), cefixime (cfm), ciprofloxacine (cip), and ceftriaxone (cro).

### 3 Results

We applied the method to 616 pneumococcal genomes from a carriage study in Massachusetts children (Nicholas J Croucher et al. 2013; Nicholas J. Croucher et al. 2015) and 1102 clinical gonococcal isolates collected from 2000 to 2013 by the Centers for Disease Control and Prevention's Gonococcal Isolate Surveillance Project (GISP) (Grad et al. 2016). For all isolates, we inferred the resistance categories as described in the main manuscript, including the ancestral state responstruction step. As lineages we used the sequenced clusters computed using Bayesian Analysis of Population Structure (BAPS) (Cheng et al. 2013).

Lineages of *S. pneumoniae* are predictive for benzylpenicillin, ceftriaxone, trimethoprim-sulfamethoxazole, erythromycin, and tetracycline resistance with AUC ranging from 0.90 to 0.97 (Figure 2). In *N. gonorrhoeae* ciprofloxacin, ceftriaxone, and cefixime attained comparably large AUCs (from 0.93 to 0.98) whereas azithromycin demonstrated lower association (AUC 0.80) (Figure 3).
