## Supplementary figures and images for "Rapid heuristic inference of antibiotic resistance and susceptibility by genomic neighbor typing"

### Figure 1

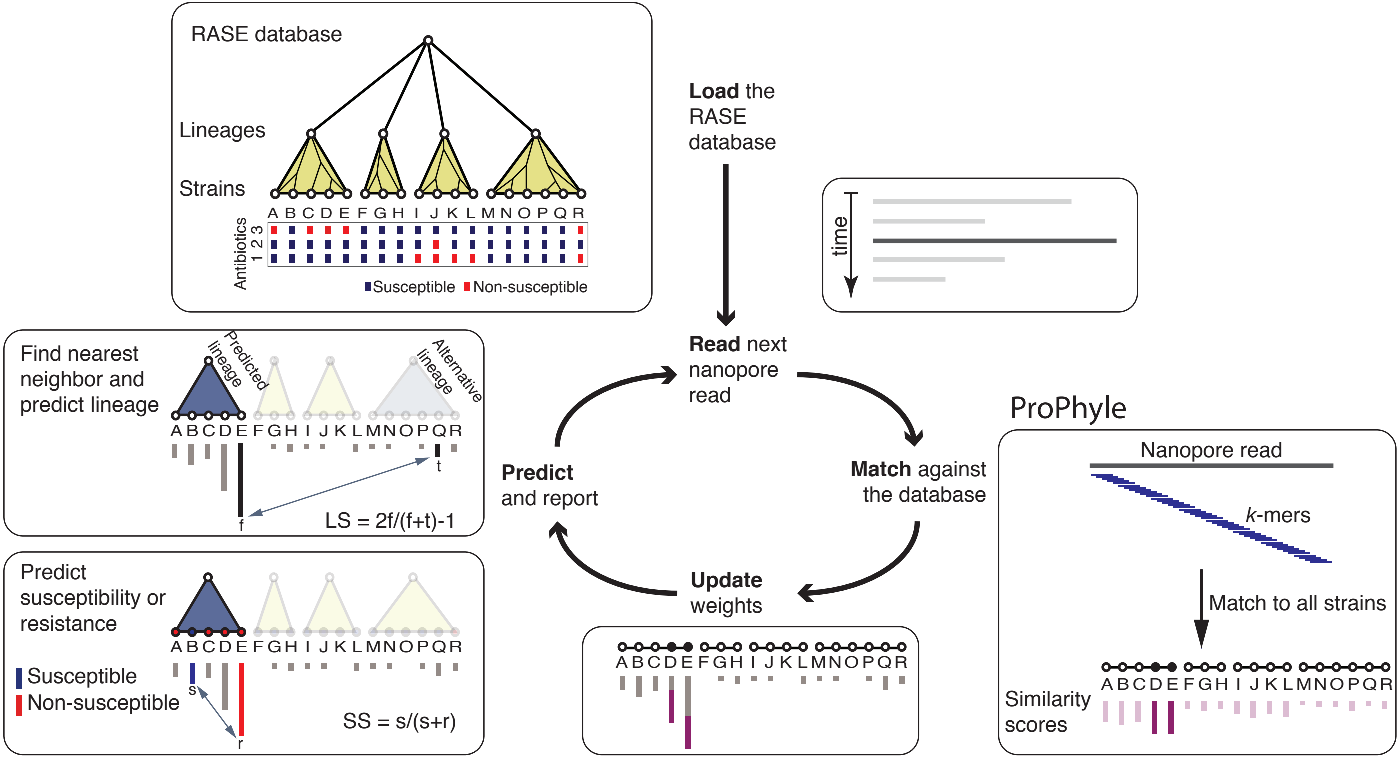

### Figure 1

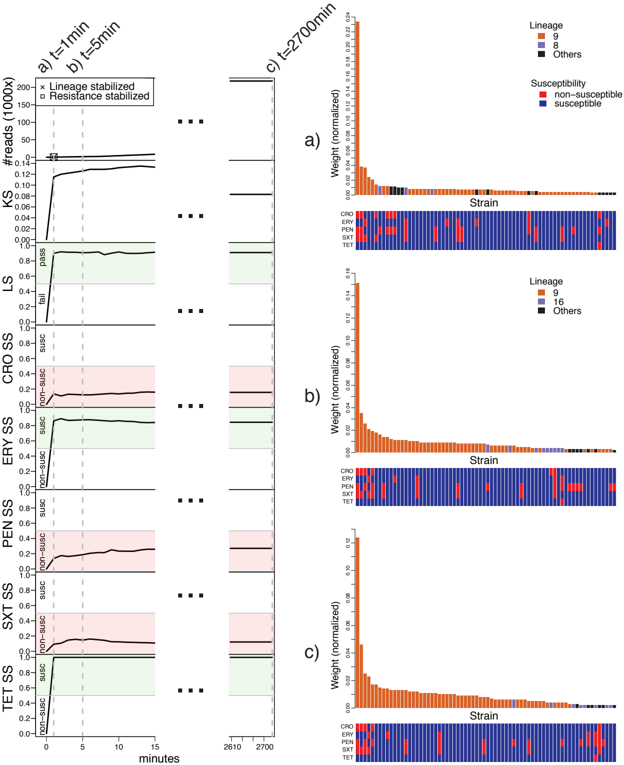

### Supplementary Figure 1

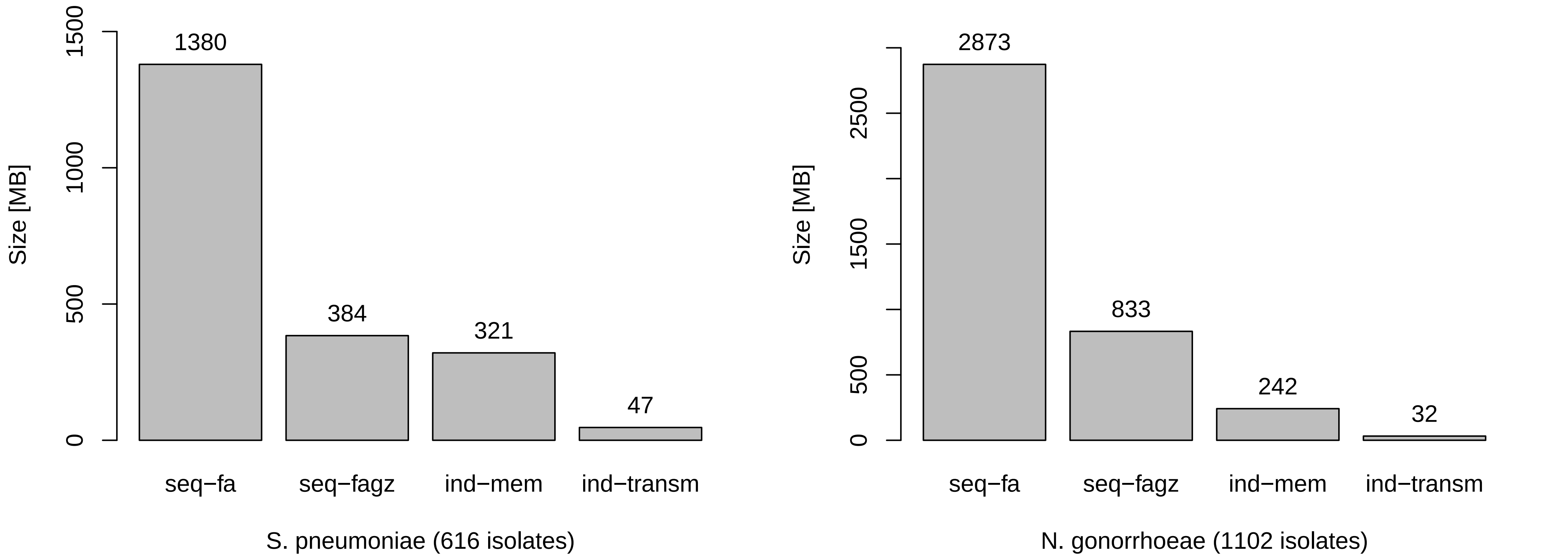

### Supplementary Figure 2

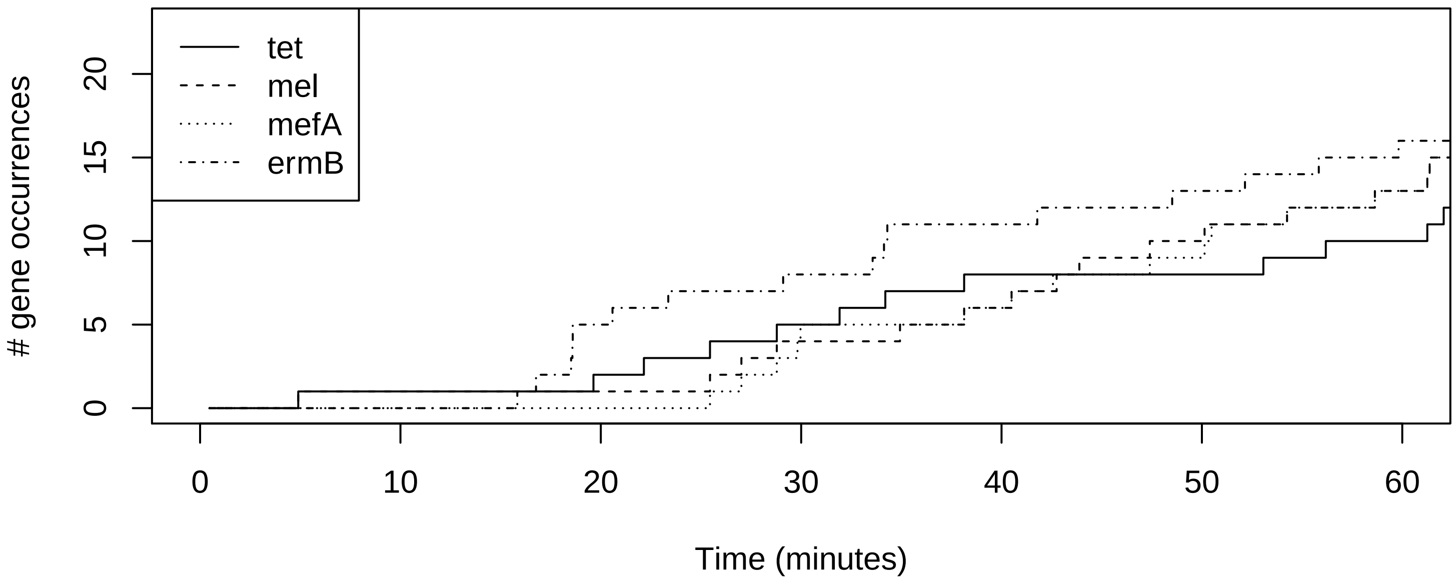

### Supplementary Figure 3

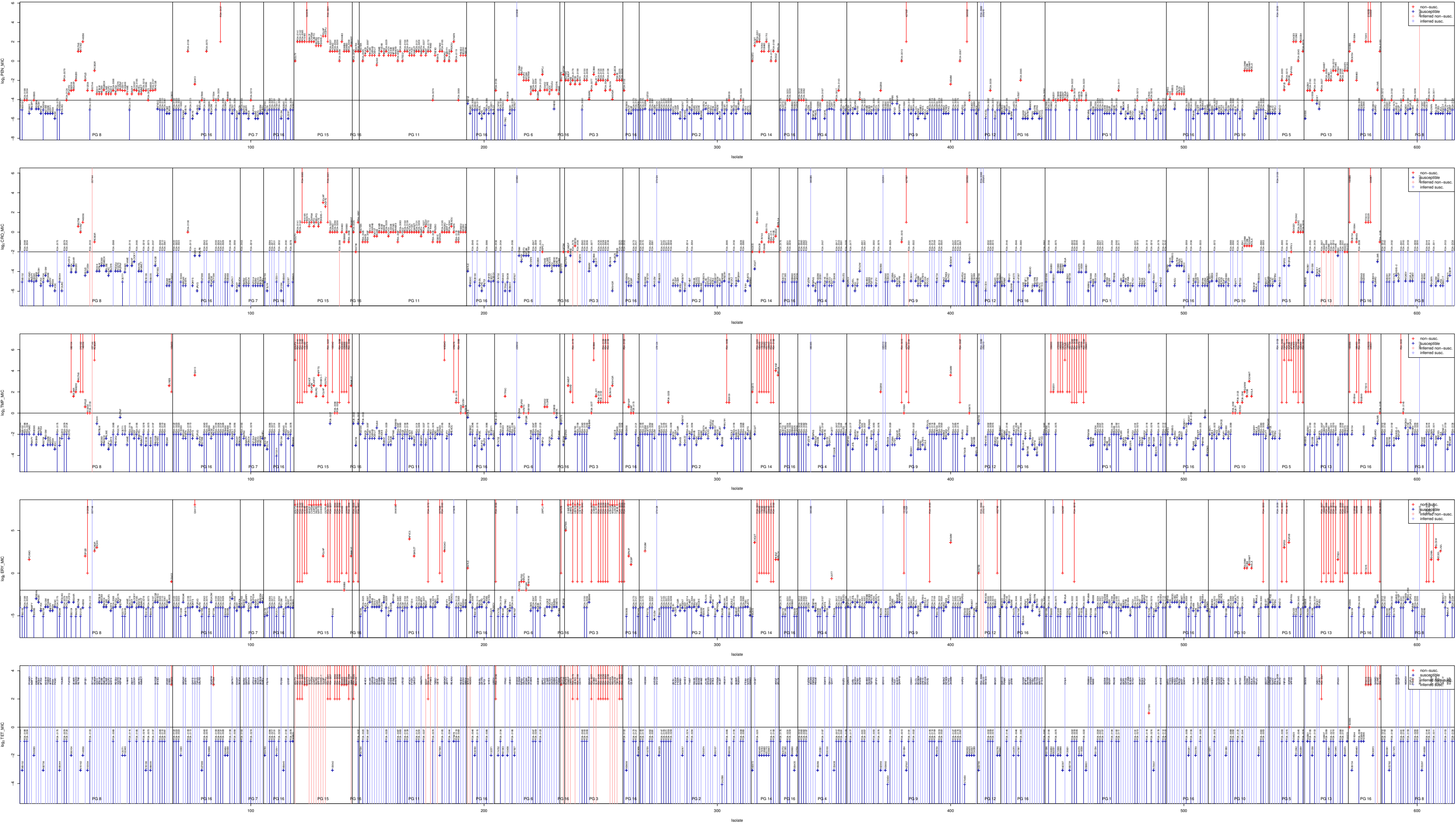

### Supplementary Figure 4

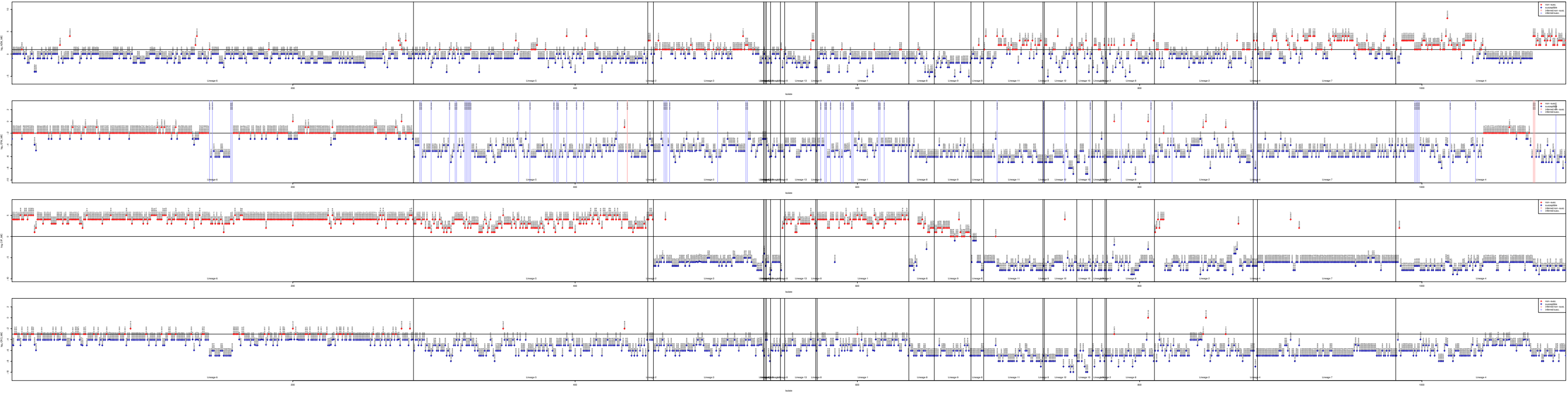

### Supplementary Figure 5

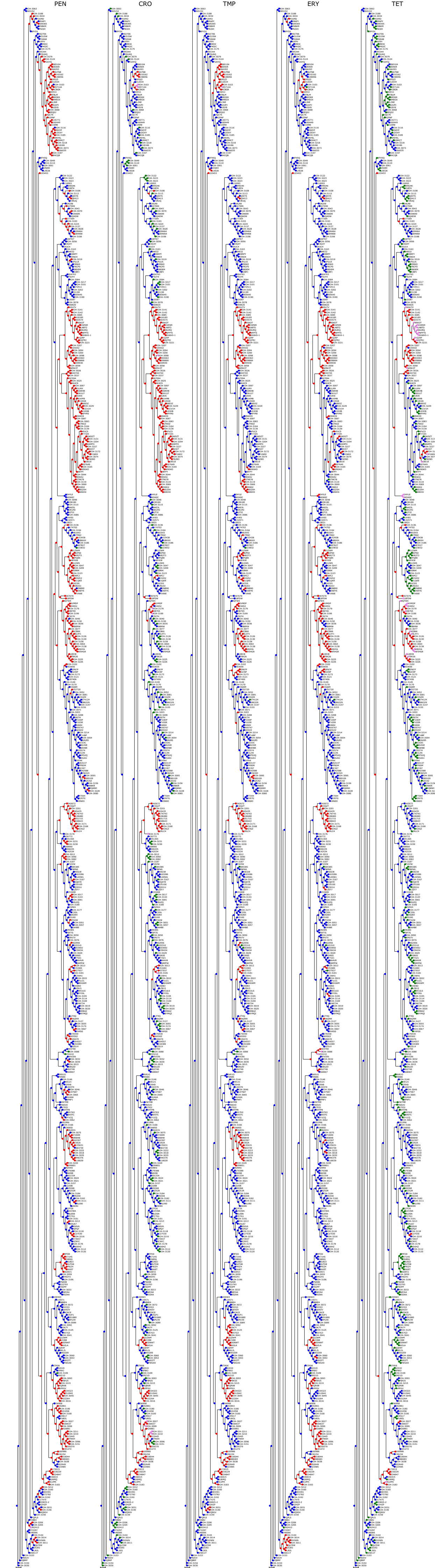

### Supplementary Figure 6

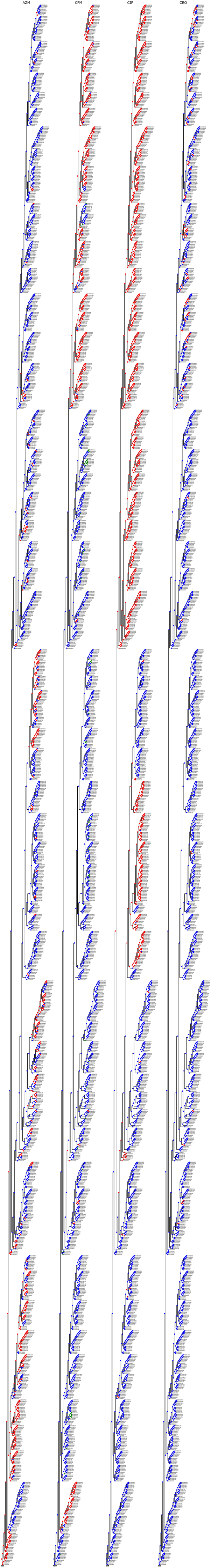

### Supplementary Figure 7

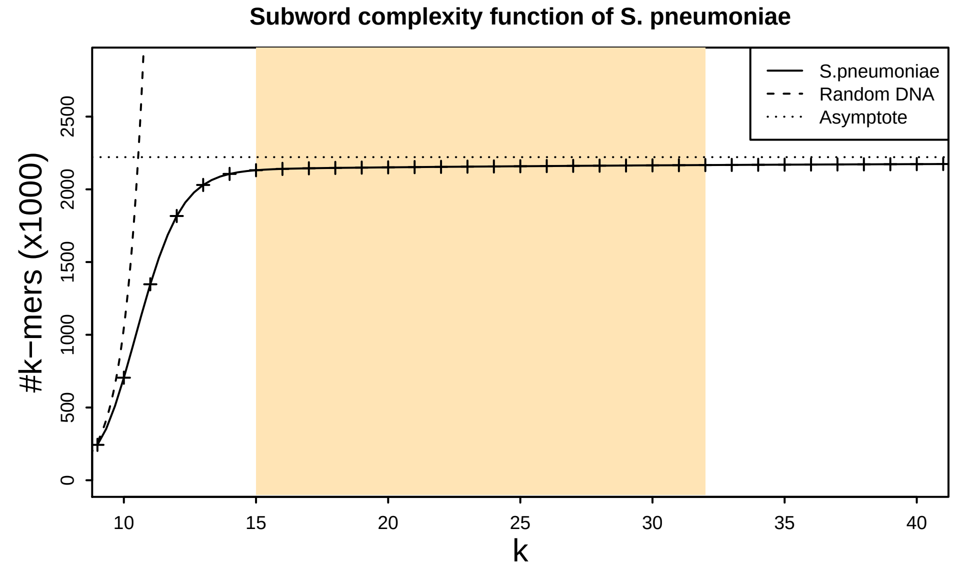

### Supplementary Figure 8

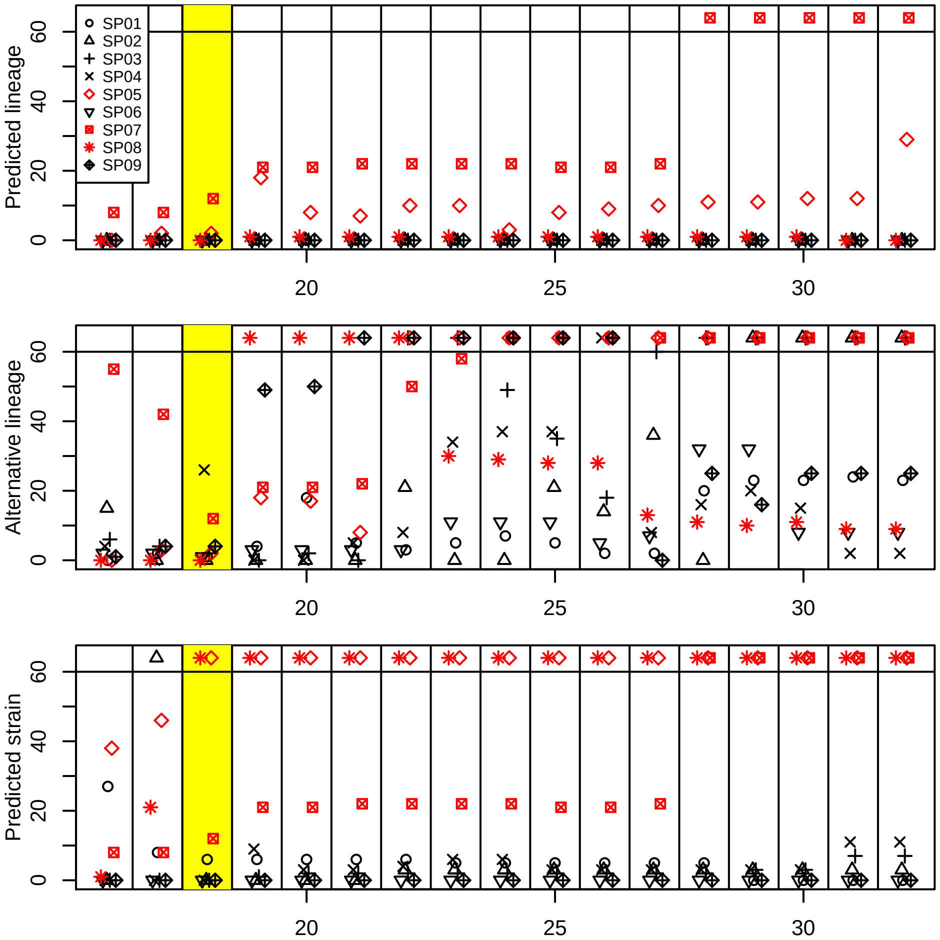
