## Supplementary Table 1 for "Rapid heuristic inference of antibiotic resistance and susceptibility by genomic neighbor typing"

### WHO lineages

| Sample | Region | Lineage confidently detected | Matched k-mers | Antibiogram AZM | | Antibiogram CFM | | Antibiogram CIP | | Antibiogram CRO | | MLST match |
|---|---|---|---|---|---|---|---|---|---|---|---|---|
| | | | | Actual | Best match | Actual | Best match | Actual | Best match | Actual | Best match | |
| WHO F (2008) | Canada | no | 17% | S | S! | S | S | S | S | S | S | OoD |
| WHO G (2008) | Thailand | no | 14% | S | S | S | S | S | R | S | S | OoD |
| WHO K (2008) | Japan | yes | 20% | S | S | R | R | R | R | S | S | yes |
| WHO L (2008) | Asia | yes | 20% | S | S | S | R | R | S | R | R | OoD |
| WHO M (2008) | Philippines | yes | 21% | S | R | S | S | R | S | S | S | yes |
| WHO N (2008) | Australia | no | 19% | S | S | S | S | R | R | S | S | OoD |
| WHO O (2008) | Canada | yes | 20% | S | S | S | S | S | S | S | S | yes |
| WHO P (2008) | USA | yes | 19% | R | R | S | S | S | S | S | S | OoD |
| WHO U (2016) | Sweden | yes | 20% | R | R | S | S | S | S | S | S | yes |
| WHO V (2016) | Sweden | yes | 19% | R | S | S | S | R | R | S | S | yes |
| WHO W (2016) | Hong Kong | yes | 20% | S | S | R | R | R | R | S | S | yes |
| WHO X (2016) | Japan | yes | 21% | S | S | R | R | R | R | R | S | yes |
| WHO Y (2016) | France | no | 18% | S | S | R | S | R | R | R | S | yes |
| WHO Z (2016) | Australia | yes | 19% | S | S | R | R | R | R | R | S | yes |
