## Supplementary material for "Rapid heuristic inference of antibiotic resistance and susceptibility by genomic neighbor typing": Table 1

### a) Database isolates

| Sample | Lineage confidently detected | Matched k-mers | Serotype | | Antibiogram CRO | | Antibiogram ERY | | Antibiogram PEN | | Antibiogram SXT | | Antibiogram TET | | MLST match | CC match |
|---|---|---|---|---|---|---|---|---|---|---|---|---|---|---|---|---|
| | | | Actual | Best match | Actual | Best match | Actual | Best match | Actual | Best match | Actual | Best match | Actual | Best match | | |
| SP01 | yes | 16% | 11D | 11D | S | S | S | S | S | S | S | S | **S[1]** | **S[1]** | Yes | Yes |
| SP02 | yes | 9.6% | 19A | 19A | R | R | R | R | R | R | R | R | **R[2]** | **R[2]** | Yes | Yes |

### b) Non-database isolates

| Sample | Lineage confidently detected | Matched k-mers | Serotype | | Antibiogram CRO | | Antibiogram ERY | | Antibiogram PEN | | Antibiogram SXT | | Antibiogram TET | | MLST match | CC match |
|---|---|---|---|---|---|---|---|---|---|---|---|---|---|---|---|---|
| | | | Actual | Best match | Actual | Best match | Actual | Best match | Actual | Best match | Actual | Best match | Actual | Best match | | |
| SP03 | yes | 3.1% | 23F | 23F | R | R | R | **S[3]** | R | R | R | R | S | S | OoD | Yes |
| SP04 | yes | 12% | 19A | 19A | R | R | R | R | R | R | R | R | R | **R[4]** | OoD | Yes |
| SP05 | no | 1.8% | 19F | 19F | R | R | R | R! | R | R | R | R! | R | R! | OoD | Yes |
| SP06 | yes | 8.3% | 23F | 23F | R | R | R | **S[3]** | R | R | R | R | S | S | OoD | Yes |

### c) Metagenomes

| Sample | Lineage confidently detected | SP | Matched k-mers | Antibiogram ERY | | Antibiogram PEN | | Antibiogram TET | |
|---|---|---|---|---|---|---|---|---|---|
| | | | | Actual | Best match | Actual | Best match | Actual | Best match |
| SP07 | no | 2.3% | 0.2% | NA | S | S | S | R | **S[5]** |
| SP08 | no | 2.5% | 0.9% | S | S | S | S! | S | **S[6]** |
| SP09 | no | 4.0% | 1.2% | NA | S | S | S | S | **S[7]** |
| SP10 | yes | 21% | 5.2% | R | R | R | R | R | **R[8]** |
| SP11 | yes | 70% | 14% | R | R | R | R | R | **R[8]** |
| SP12 | yes | 86% | 17% | S | S | S | S | R | **S[5]** |

**Legend**

Correct prediction
Incorrect prediction
Cannot be evaluated

| | |
|---|---|
| S | Susceptible |
| R | Non-susceptible |
| ! | Low confidence call |
| NA | Not available |
| OoD | Out-of-database |
| (...) | ID of a retested sample |
| SP | Fraction of S. pneumoniae reads |
