## Supplementary material for "Rapid heuristic inference of antibiotic resistance and susceptibility by genomic neighbor typing": Table 2

### a) Database isolates

| Sample | Lineage confidently detected | Matched k-mers | Antibiogram AZM | | Antibiogram CFM | | Antibiogram CIP | | Antibiogram CRO | | MLST match |
|---|---|---|---|---|---|---|---|---|---|---|---|
| | | | Actual | Best match | Actual | Best match | Actual | Best match | Actual | Best match | |
| GC01 | yes | 27% | S | S | S | S | S | S | S | S | Yes |
| GC02 | yes | 27% | S | S | R | R! | S | S | R | R! | Yes |
| GC03 | yes | 33% | S | S | R | S! | S | S | R | S! | Yes |
| GC04 | yes | 21% | S | S | R | R | R | R | R | S | Yes |
| GC05 | yes | 7% | R | R | S | S | S | S | S | S | Yes |

### b) Clinical isolates

| Sample | Lineage confidently detected | Matched k-mers | Antibiogram AZM | | Antibiogram CFM | | Antibiogram CIP | | Antibiogram CRO | |
|---|---|---|---|---|---|---|---|---|---|---|
| | | | Actual | Best match | Actual | Best match | Actual | Best match | Actual | Best match |
| GC06 | yes | 19% | S | S | R | R | R | R | S | S |
| GC07 | no | 20% | S | S | S | S | R | R | S | S |
| GC08 | no | 19% | S | S | S | S | R | R | S | S |
| GC09 | no | 18% | S | S | S | S | S | S | S | S |
| GC10 | no | 20% | S | S | S | S | R | R | S | S |
| GC11 | no | 20% | S | S | S | S | R | R | S | S |
| GC12 | no | 20% | S | S | S | S | R | R | S | S |
| GC13 | yes | 20% | S | S | S | S | R | R | S | S |
| GC14 | yes | 19% | S | S | S | S | R | R | S | S |
| GC15 | yes | 19% | R | S! | S | S | S | S | S | S |
| GC16 | no | 18% | S | S | S | S! | R | R | S | S! |
| GC17 | no | 19% | S | S | S | S! | R | R | S | S! |
| GC18 | no | 20% | S | S | S | S | R | R | S | S |
| GC19 | yes | 18% | S | S | S | S | R | R | S | S |
